# Oxysterol-liver X receptor signaling mediates CYFIP1 regulation of cortical neurogenesis

**DOI:** 10.1101/2023.06.23.546272

**Authors:** Daniel Cabezas De La Fuente, Claudia Tamburini, Emily Stonelake, William J Griffiths, Jeremy Hall, Michael J. Owen, David E. J. Linden, Andrew Pocklington, Yuqin Wang, Meng Li

## Abstract

Dysregulation in neural progenitor proliferation and neuronal differentiation has been increasingly recognised as a common pathology in neural cells harboring genetic risks to neuropsychiatric and neurodevelopmental disorders, yet the underlying molecular mechanisms remains largely unknown. Deletions and duplications of the 15q11.2 region containing the CYFIP1 gene have been associated with autism and schizophrenia. Using patient-derived iPSCs carrying 15q11.2 deletion and genetically manipulated hESCs with CYFIP1 gain- and loss-of-function (GoF and LoF), we show that 15q11.2 deletion and CYFIP1-LoF leads to premature neuronal differentiation while CYFIP1-GoF favours neural progenitor maintenance. We identified cholesterol biosynthesis and metabolism as a biological process disturbed by CYFIP1 dosage change, leading to altered neuro-oxysterol profiles. 24S,25-epoxycholesterol, which was decreased in CYFIP1-GoF and increased in CYFIP1-LoF and 15q11.2del neural cells, can mimic the 15q11.2del and CYFIP1-LoF phenotype by promoting cortical neuronal differentiation and restore the impaired neuronal differentiation of CYFIP1-GoF neural progenitors. Moreover, the neurogenic activity of 24S,25-epoxycholesterol is lost following genetic deletion of the brain expressed isoform of the liver X receptor LXRb while compound deletion of LXRb in CYFIP1-/- background rescued their premature neurogenesis. This work delineates LXR mediated oxysterol regulation of neurogenesis as a novel pathological mechanism in neural cells carrying 15q11.2CNV and provides a potential target for therapeutic strategies for genetic disorders associated with this risk locus.

## Introduction

Neurodevelopmental disorders, such as autism and schizophrenia, are believed to originate during early brain development with shared genetic etiology^1^. Copy number variations (CNVs) at the chromosomal locus 15q11.2 (BP1-BP2) have been found by several genome wide association studies to be significantly linked to an increased risk of developing these disorders^2–6^. In addition to psychiatric features, the clinical phenotype of 15q11.2 CNVs carriers frequently includes other traits, such as developmental delay, seizures and motor coordination problems^7^. The 15q11.2 region spans approximately 500 kb and contains four genes encoding CYFIP1, NIPA1, NIPA2 and TUBGCP5^8^. CYFIP1 (cytoplasmic FMR1 interacting protein), is considered the strongest candidate underlying the link between 15q11.2 CNVs and psychiatric illness due to its functional association with the fragile X mental retardation protein (FMRP)^9,10^. In addition to regulating mRNA translation together with FMRP, CYFIP1 is also part of the WAVE complex that controls actin polymerisation^11^. CYFIP1 is enriched in dendritic spines, and alterations in its level in rodent primary cultures and genetic mouse models affect dendritic morphology and synaptic function^12–15^, disrupt adult neurogenesis^16, 17^, and cause white matter changes^18, 19^.

While previous studies into CYFIP1 mostly focused on the adult brain, transcript levels of CYFIP1 are highest in the first trimester of human brain development (http://hbatlas.org/pages/hbtd) and in situ hybridisation of fetal mouse brain shows Cyfip1 RNA concentrated in the ventricular zone and sub-ventricular zone (http://www.eurexpress.org). Consistent with its early expression profile, shRNA knock down of CYFIP1 in hESC-derived neural progenitors disrupts their polarity, while electroporation of shRNA-Cyfip1 into the developing mouse cortex affects cortical progenitor migration^20^. Neuroepithelial polarity and progenitor migration are intricately associated with neural progenitor cell (NPC) division and neurogenesis. The above findings thus raise the question whether CYFIP1 deficiency leads to aberrant progenitor proliferation and neuronal differentiation. Moreover, whether an increase in CYFIP1 protein level, as in 15q11.2 duplication, affects cortical neural induction and neuronal function remains unknown.

Using transgene expression and CRISPR/Cas9 assisted genome-editing of *CYFIP1* in human embryonic stem cells (hESC) and patient-derived induced pluripotent stem cells (iPSC) harbouring 15q11.2 chromosomal deletions (15q11.2del iPSCs), we find that gain- and loss-of CYFIP1 disturb cortical NPC proliferation and differentiation with opposing directions of effect. Defects caused by CYFIP1 deficiency were mirrored by 15q11.2del iPSC-derived neural cells. Moreover, we provide unexpected evidence that CYFIP1-dependent neurogenesis is regulated through controlling biosynthesis and metabolism of oxysterols, which directly affects neurogenic activity of NPCs via the liver X receptor (LXR) signalling. Thus, our study identifies CYFIP1 as a new regulator for cortical neurogenesis and provides novel mechanistic insight into the causes of the neurodevelopmental impairment in 15q11.2 CNV carriers and potential therapeutic targets^21^.

## Results

### Gain and loss of CYFIP1 led to altered neuronal differentiation kinetics

To mimic CYFIP1 dosage change in 15q11.2del and 15q11.2dup we generated stable, clonal lines of hESCs that either deficient in CYFIP1 expression (CYFIP1 loss-of-function, CYFIP1-LoF) or carry a CYFIP1 transgene (CYFIP1-GoF). The CYFIP1-LoF lines were generated by CRISPR/Cas9 assisted genome editing in the iCas9 background^22^ while the CYFIP1-GoF cells are of H7 origin and ubiquitously express human CYFIP1 driven by the CAG promoter (for details see supplemental methods and results, Figure S1A, E-F). The CYFIP1-LoF lines used in this study showed ∼5% (clone #31) and 40% (clone #41) CYFP1 protein level of their isogenic wildtype control cells, respectively (Figure S1G), while the CYFIP1-GoF lines (#3 and #5) both exhibited approximately two-fold increase of CYFIP1 (Figure S1B-D).

*CYFIP1* transcript levels are highest in the first trimester of human telencephalic development (http://hbatlas.org/pages/hbtd). We therefore investigated the effect of CYFIP1 dosage change on cortical neural differentiation of CYFIP1-LoF and CYFIP1-GoF hESCs (Figure 1A). At day 15 (d15), the CYFIP1-GoF and CYFIP1-LoF cultures contained a similar number of FOXG1^+^ and PAX6^+^ forebrain progenitors compared to their respective isogenic control cultures (Figure S2), suggesting a normal cortical neural induction from human pluripotent stem cells (hPSCs). However, at d35 when neurons are actively produced, we detected a marked reduction of NeuN^+^ neurons in CYFIP1-GoF cultures compared to the isogenic controls (Figure 1B-C). The CYFIP1-LoF cultures showed the opposite phenomenon with a significant increase of NeuN^+^ cells compared to their control cultures (Figure 1C, 1D).

**Figure 1.**
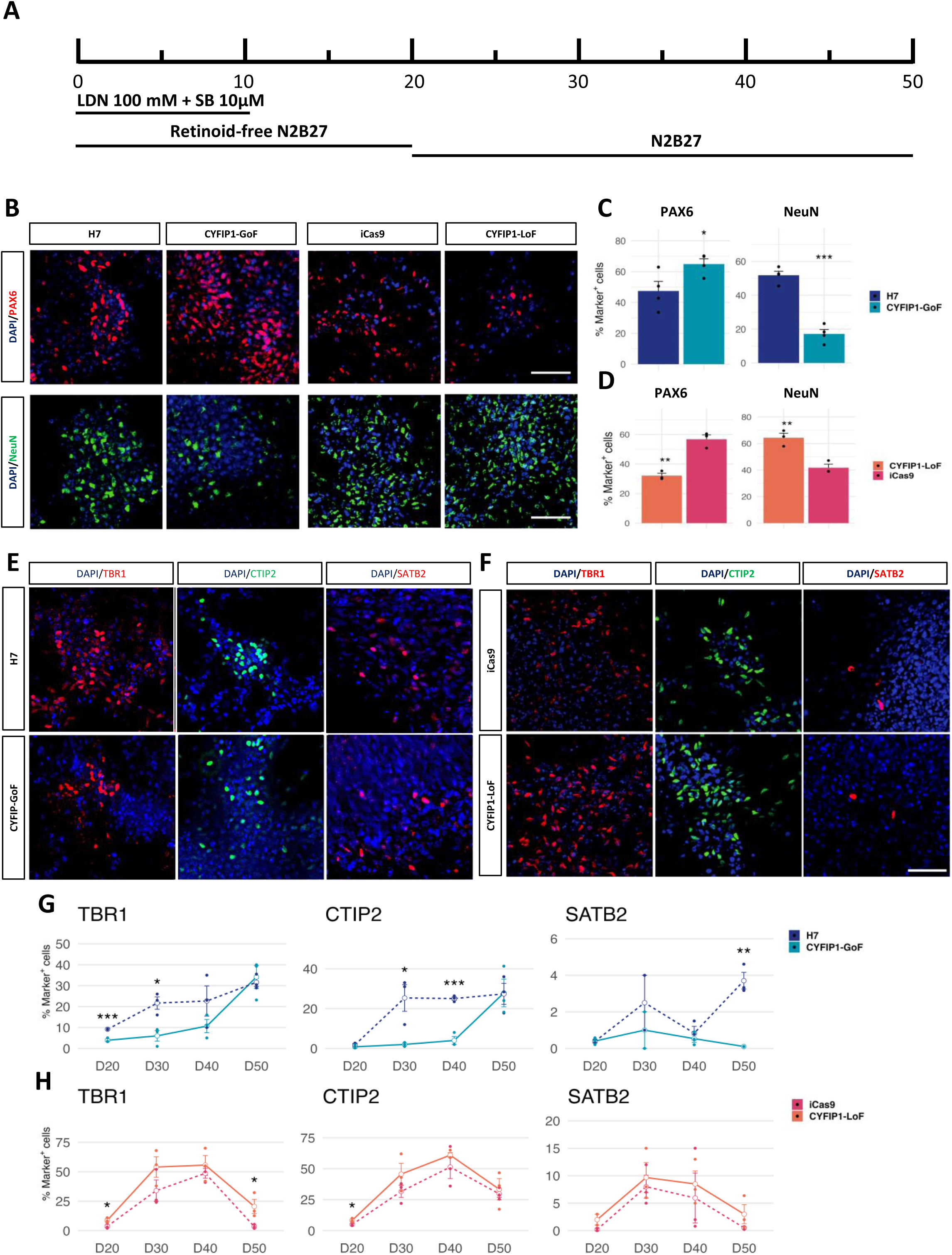
Gain- and loss-of CYFIP1 affect temporal kinetics of neuronal differentiation. A) Outline of the cortical differentiation paradigm. B) Immunostaining for PAX6 (red) and NeuN (green) for CYFIP1-GoF, CYFIP1-LoF and their respective isogenic controls at d35 of differentiation. C-D) Quantification of PAX6^+^ and NeuN^+^ cells shown in B. E-F) Day 20-50 cortical cultures were immunostained for cortical neuronal markers TBR1 (red), CTIP2 (green) and SATB2 (red), shown are representative images of day 30 staining. G-H) Quantification of TBR1^+^, CTIP2^+^ and SATB2^+^ of day 20 to day 50 cortical cultures, respectively. Data presented are mean±s.e.m. from at least three biological replicates, groups were compared by Student t-test between the CYFIP1-manipulated vs isogenic control at each time point (*p<0.05; **p<0.01; ***p<0.001). Nuclei were counterstained with DAPI (blue). Scale bars: 50 µm.

We next broadened the analysis time window from d20 till d50 of differentiation cultures and examined typical markers representing different layers of cortical neurons. CYFIP1-GoF cultures contained a lower percentage of TBR1^+^ and CTIP2^+^ cells compared to their respective isogenic controls between d20 and d40 but caught up by d50 (Figure 1 E, G). Conversely, the percentage of these neuronal subtypes were generally higher in the CYFIP1-LoF population than the isogenic controls (Figure 1F, H). SATB2 is expressed in upper layer and a subset of CTIP2^+^ cortical neurons^23^. SATB2^+^ neurons largely mirrored those for deep layer markers (direction of effect is the same), but only reached statistical significance for CYFIP1-GoF at d50. Together these experiments showed that elevated levels of CYFIP1 led to a delayed neuronal production while CYFIP1-deficiency results in precocious neuron differentiation.

### CYFIP1 dosage change affects neurogenesis rate of cortical progenitors

Given the altered neuronal differentiation kinetics observed above, we next carried out a pulse-chase EdU incorporation assay at d30 followed by double EdU/NeuN antibody staining five days later to determine the relative neuronal birth rate by CYFIP1-GoF and -LoF neural progenitors (Figure 2). We found that the CYFIP1-GoF cultures contained fewer EdU retaining NeuN^+^ neurons than the isogenic control cultures (Figure 2A-B). In contrast, more EdU-retaining NeuN^+^ neurons in CYFIP1-LoF cultures than in the respective controls (Figure 2A-C), suggesting that CYFIP1 levels regulate the rate of neurogenesis in these cortical progenitors.

**Figure 2.**
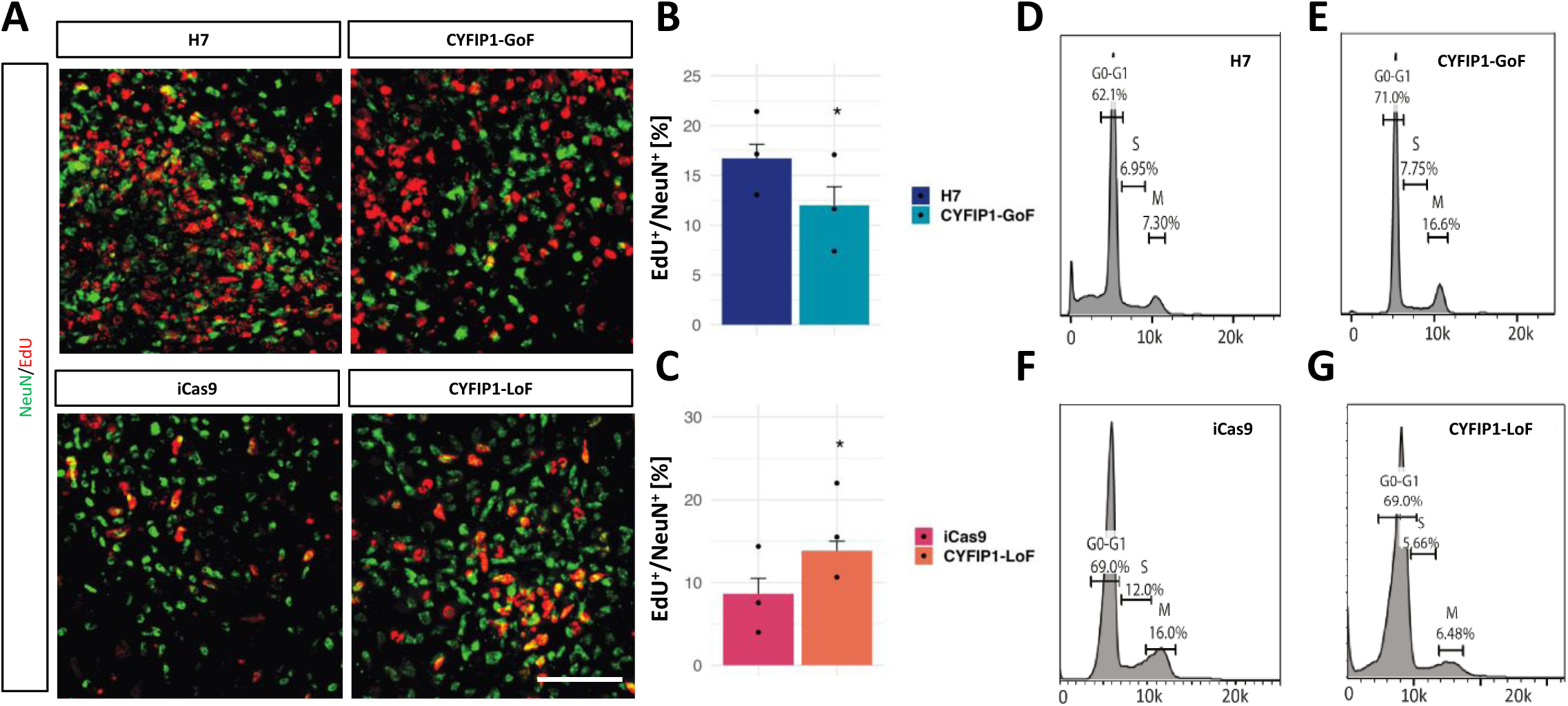
CYFIP1 dosage change affects NPC cell cycle profile. A) Representative images of cells labelled with EdU (red) and NeuN (green) at d35. D30 cultures were incubated with EdU for 2 hours and co-immunostained 5 days later for EdU and NeuN. B-C) Quantification of EdU retaining cells in NeuN^+^ neurons. D-G) Histograms of a flow cytometry-based cell cycle analysis on day 30 cultures. Data presented are mean±s.e.m. from three biological replicates. Student t-test, *p<0.05. Scale bars: 50 µm.

Consistent with the above findings, flow cytometry-based cell cycle analysis revealed a higher percentage of M phase progenitors in CYFIP1-GoF cultures than the controls (Figure 2D, E). In contrast, a reduced number of S and M phase cells were detected in the CYFIP1-LoF cultures compared to the isogenic controls, indicating that a higher proportion of CYFIP1-LoF progenitors had exited the cell cycle (Figure 2F, G). This change in cell cycle profile of CYFIP1-LoF cultures was also associated with a reduction in the number of cells positive for the proliferation marker Ki67 and a concurrent increase in cells expressing the cell cycle regulator P27kip1 (Figure S3), that can promote neuronal differentiation in the cerebral cortex^24^. These observations suggest that a tight regulation of CYFIP1 level is essential for normal cortical neural progenitor cell division and neurogenesis.

### Genome-wide transcriptome studies confirms altered neural developmental kinetics and reveals disrupted cholesterol biosynthesis and metabolism

In order to gain a deeper understanding of the cellular mechanisms and pathways disrupted by CYFIP1 dosage change, we performed RNA-sequencing (RNAseq) of CYFIP1-GoF and CYFIP1-LoF and their respective parental control cells at three time points of cortical neural differentiation, which represent an early NPC (NPC1, d10), peak NPC and onset of neurogenesis (NPC2, d20) and neuronal rich (Neuron, d40) stages, respectively (Figure 3A). Principal component analysis (PCA) revealed a clear differentiation stage (PC1) and genotype (PC2) dependent separation in both the CYFYP1GoF and CYFIP1-LoF datasets (Figure S4A, B). Transcription factors and signalling molecules important for cortical development and progenitor proliferation, such as PAX6, EMX1, LHX2, frizzled receptors and dopachrome tautomerase (DCT)^25^ were amongst the differentially expressed genes (DEGs) between cells with CYFIP1 dosage change and their isogenic controls(Figure 3B-C, Table S1-2). Interestingly, many DEGs showed opposite direction of change in CYFIP1-GoF and CYFIP1-LoF samples. Significant changes were also detected for genes associated with neuronal differentiation and synapse formation, such as CDH1, doublecortin (DCX), Synaptotagmins (SYT1 and SYT2)^26^ and DLG4 (PSD95) (Figure 3B-C).

**Figure 3.**
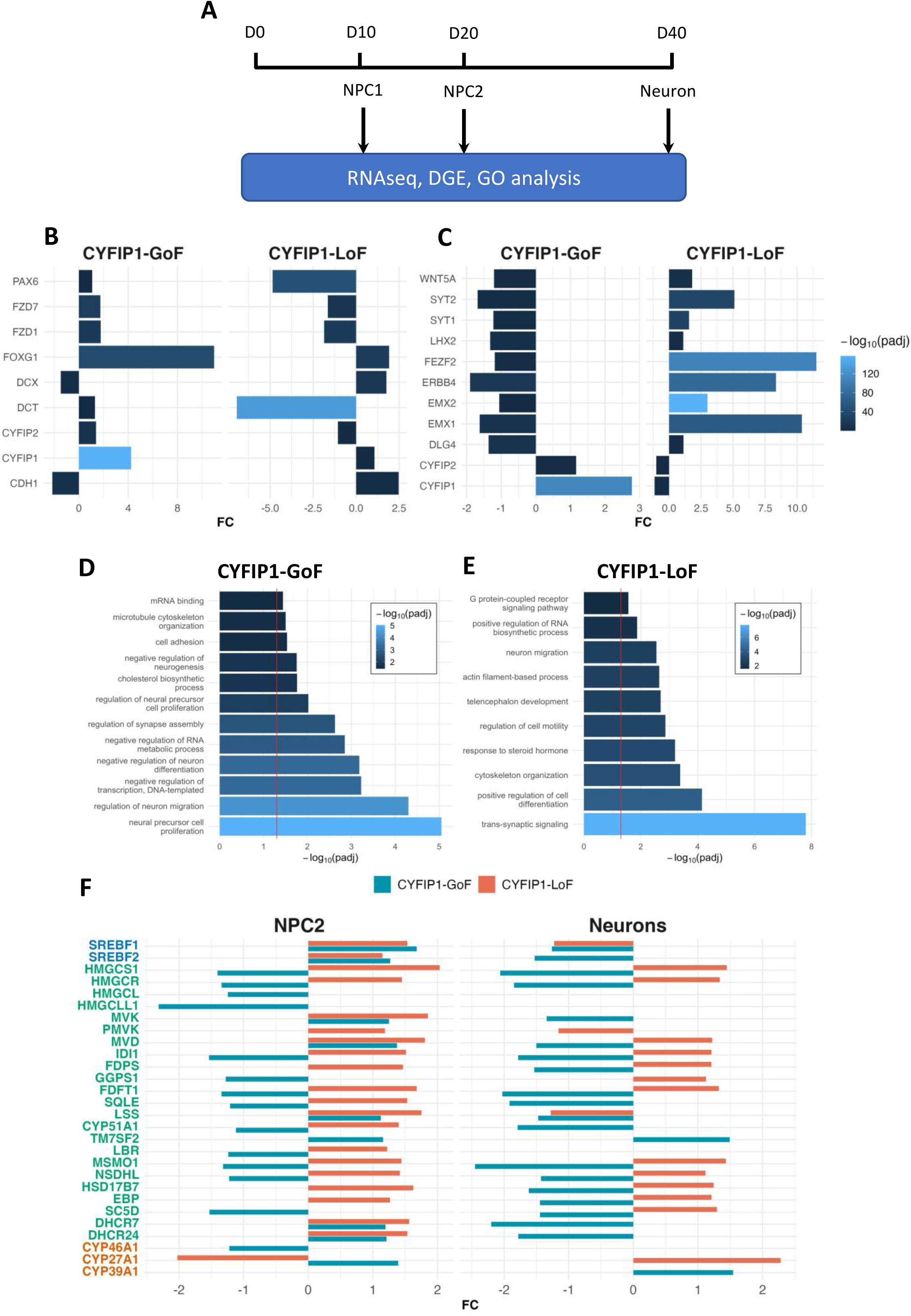
Genome-wide transcriptomic analysis of CYFIP1-GoF and CYFIP-LoF neural cells. A) Schematic illustration of the RNAseq experimental flow. B-C) Bar graphs showing examples of DEGs related to neural progenitors and neuronal differentiation at NPC2 (B) and neuronal stage (C), respectively. Bar height represents fold change (FC) the shade of color represents the FDR adjusted p value. D-E) Gene ontologies (GO) enriched in the DEG datasets associated with CYFIP1-GoF (D) and CYFIP1-LoF (E), compared to the respective control samples for all developmental stages analysed. Bar height represents the -log10 of the Benjamini-Hochberg adjusted P value, red line shows the threshold of significance (i.e. padj < 0.05). F) Bar graphs showing DEGs involved in cholesterol biosynthesis (green) and metabolism (red) and the master transcription factors for fatty acid and cholesterol biosynthesis (blue) at NPC2 and neuronal stages.

To reveal CYFIP1-regulated biological processes and molecular pathways, we analyzed CYFIP1-GoF and CYFIP1-LoF protein-coding DEGs separately at each developmental stage to identify enriched gene ontology (GO) terms. GO terms significantly over represented (Benjamini Hochberg-corrected P < 0.05) amongst DEGs were primarily linked to the regulation of central nervous system development (Figure 3D-E), reflecting the findings of altered differentiation kinetic of NPCs and their direction of change. For example, ‘neural precursor cell proliferation’ was amongst the most significant terms in CYFIP1-GoF samples while one of the highest in the list of GO terms associated with CYFIP1-LoF was ‘cell differentiation’. Terms relating to microtubule cytoskeleton organization and synaptic assembly were also significantly enriched, indicating that our results are in line with previously reported functions of CYFIP1 (Figure 3D-E, Table S3-4). Unexpectedly, this analysis identified the presence of significant alterations of cholesterol biosynthesis and sterol response in both CYFIP1-GoF and CYFIP1-LoF samples, as demonstrated by differential expression, mostly in contrasting directions, of genes involved in the cholesterol biosynthetic process (Figure 3F, table S1-2). Examples of which include *SREBF1* & *SREBF2*, the genes that encode the sterol regulatory element binding proteins 1 & 2 (SREBP1 & 2) required for fatty acid and cholesterol synthesis, and most of the genes for enzymes in cholesterol biosynthesis cascade.

### Convergent premature neuronal differentiation of 15q11.2del neural progenitors

CYFIP1 is one of the four genes in the 15q11.2 locus (*CYFIP1*, *NIPA1*, *NIPA2* and *TUBGCP5*). To directly compare the cellular and molecular event consequent of CYFIP1-LoF and 15q11.2del, we derived iPSCs from two individuals carrying 15q11.2 deletions (referred to as EA8 and EA62, respectively) and control iPSCs from two unaffected controls (CTRL-iPSC). Following the same cortical differentiation paradigm, we observed evident reduction of CYFIP1 protein level in both NPCs and neurons derived from iPSCs of the two deletion carriers (two clones each) compared to respective control cell types (Figure S4C-D). Interestingly, NIPA2 expression was only detected in NPCs and not in neurons of either the control iPSC- or 15q11.2del iPSC-derivatives. 15q11.2del NPCs had a lower level of NIPA2 compared to the controls (Figure S4C-D). TUBGCP5 on the other hand was detected in both NPC and neuronal samples and the expression level was not affected by 15q11.2 deletion (Figure S4C-D). We were not able to detect NIPA1 protein in any of the samples tested.

Similar to the observations made in CYFIP1-LoF cultures, 15q11.2del iPSC-derived cultures contained fewer PAX6^+^, Ki67^+^ cells and more P27kip1^+^ cells than the control cultures at d35 (Figure 4A-C). In contrast, both iPSC-derived 15q11.2del lines tended to produce a higher proportion of TBR1^+^ and CTIP2^+^ neurons at d30 and d40 although line variation was apparent (Figure 4D-E). These data nonetheless indicate that 15q11.2 deletion leads to premature neurogenesis, as observed in CYFIP1-LoF.

**Figure 4.**
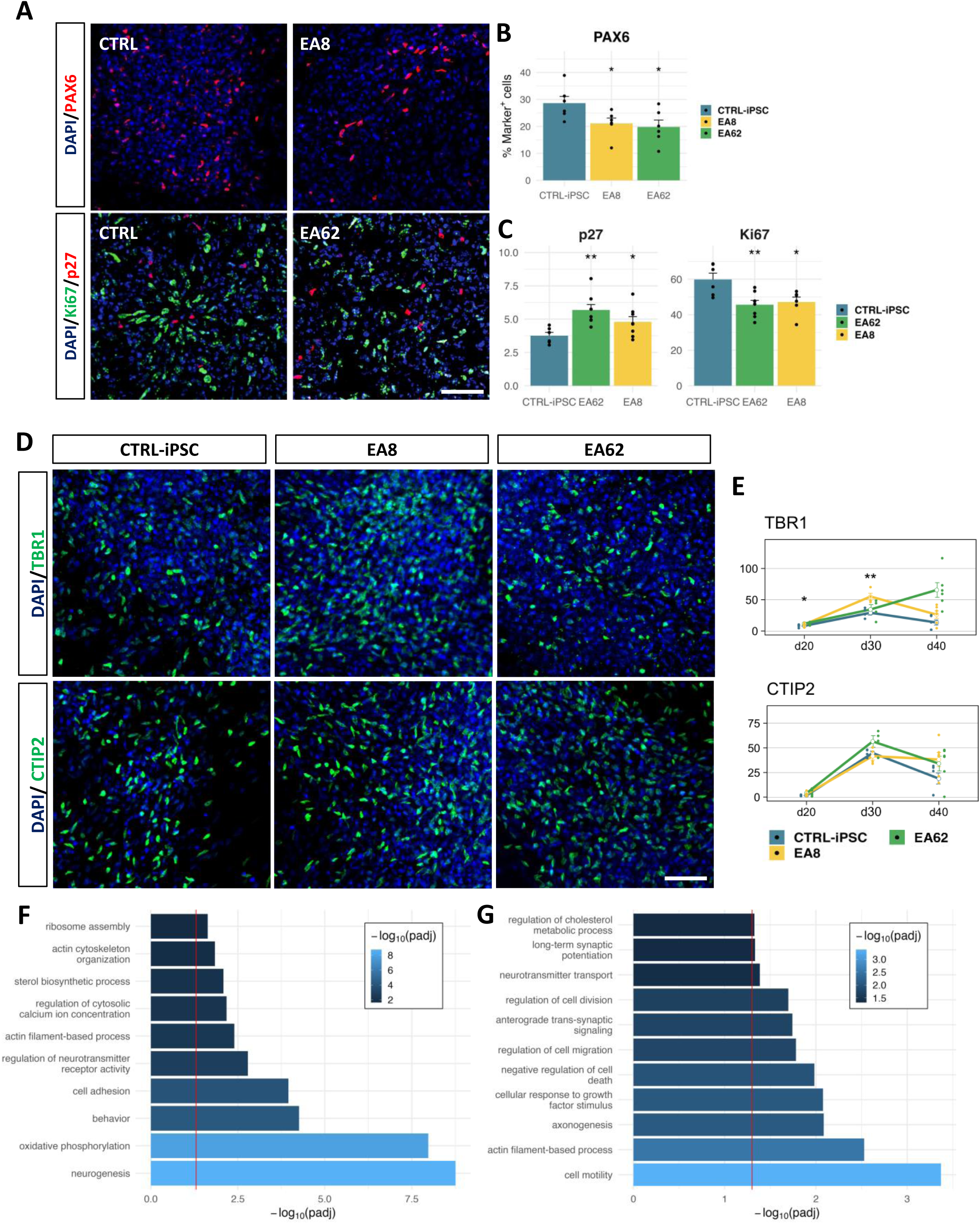
15q11.2del iPSCs display a similar neurogenesis phenotype to that of CYFIP1-LoF. A) Two clones each of EA8 and EA62 15q11.2del lines and two control iPSC lines were differentiated towards cortical fate, immunostaining was performed at d35 cultures for PAX6 (red), Ki67 (green) and p27^kip1^ (red). B-C) Quantification data for the proportion of PAX6^+^, Ki67^+^ and p27^kip1+^ cells. D) Immunostaining performed at d30 for deep layer cortical neuronal markers TBR1 and CTIP2 (green). E) Quantification data for the proportion of TBR1^+^ and CTIP2^+^ cells at d20, 30 and 40. F-G) Gene ontologies (GO) enriched in the DEG datasets associated with EA8 (F) and EA62 (G) compared to the control iPSC samples. Bar height represents the -log10 padj value, red line shows the threshold of significance (i.e. padi< 0.05). Each dot in B, C and E represent a biological replicate. Data presented are mean±s.e.m. from at least three biological replicates for each of the clonal lines. Groups were compared by Student t-test between the control iPSC and each of the 15q11.2del lines per time point (*p<0.05, ** p<0.01). Nuclei were counterstained with DAPI (blue). Scale bars: 50 µm.

To gain further insights into the cellular and molecular changes caused by 15q11.2 deletion and investigate how these changes relate to loss of CYFIP1 alone, we carried out RNAseq on d20 NPCs and d40 neurons derived from EA8 and EA62 iPSCs and a control iPSC line (Figure S4E). This experiment revealed around 2fold reduction of *CYFIP1*, *NIPA1*, *NIPA2* and *TUBGCP5* transcripts in samples from both deletion carriers (Figure S4F, Table S5-6), reflecting the effect of the 15q11.2del and proving the reliability of our transcriptomic approach. GO analysis showed a significant dysregulation of biological processes relative to neurogenesis, cell proliferation and synaptic signaling in samples from both 15q11.2del carriers, and time points analyzed (Figure 4F-G, Table S7-8). Importantly, consistent with findings in the CYFIP1-LoF neural samples, sterol and cholesterol biosynthetic and metabolic process were also amongst the significantly altered biological processess in both 15q11.2del iPSC lines (Figure 4F, G).

Together, our findings demonstrate that 15q11.2 deletion results in phenotypic and transcriptomic alterations analogous to those caused by loss of CYFIP1 alone, highlighting a causal role for CYFIP1 in the pathological manifestations associated with 15q11.2 CNVs. Crucially, our transcriptomic analysis of CYFIP1-manipulated and 15q11.2del neural cells both revealed sterol biosynthesis as a novel CYFIP1-dependent function.

### 15q11.2del and CYFIP1-GoF cells display contrasting profile changes of 24S,25-epoxycholesterol

In light of the dysregulated cholesterol biosynthesis and metabolism revealed by the transcriptomics analysis, we performed a metabolic profiling of NPCs and neurons derived from both CYFIP1-LoF and CYFIP1-GoF lines and the two 15q11.2del iPSC lines by liquid chromatography-mass spectrometry with multistage fragmentation (LC-MS^n^). This analysis indeed identified striking alterations in neuro-sterols and oxysterols (oxidized forms of cholesterol or of its precursors). Compared to the respective isogenic hESC- and control iPSC-derived NPCs and neuronal cells, both CYFIP1-LoF and 15q11.2del samples showed an increased level of 24S,25-epoxycholesterol (24S,25-EC), an oxysterol generated in the shunt pathway during cholesterol biosynthesis and also from desmosterol via oxidation by the enzyme CYP46A1. In contrast, CYFIP1-GoF cells showed a decrease of 24S,25-EC in comparison to their isogenic control cells of corresponding differentiation stage (Figure 5A).

**Figure 5.**
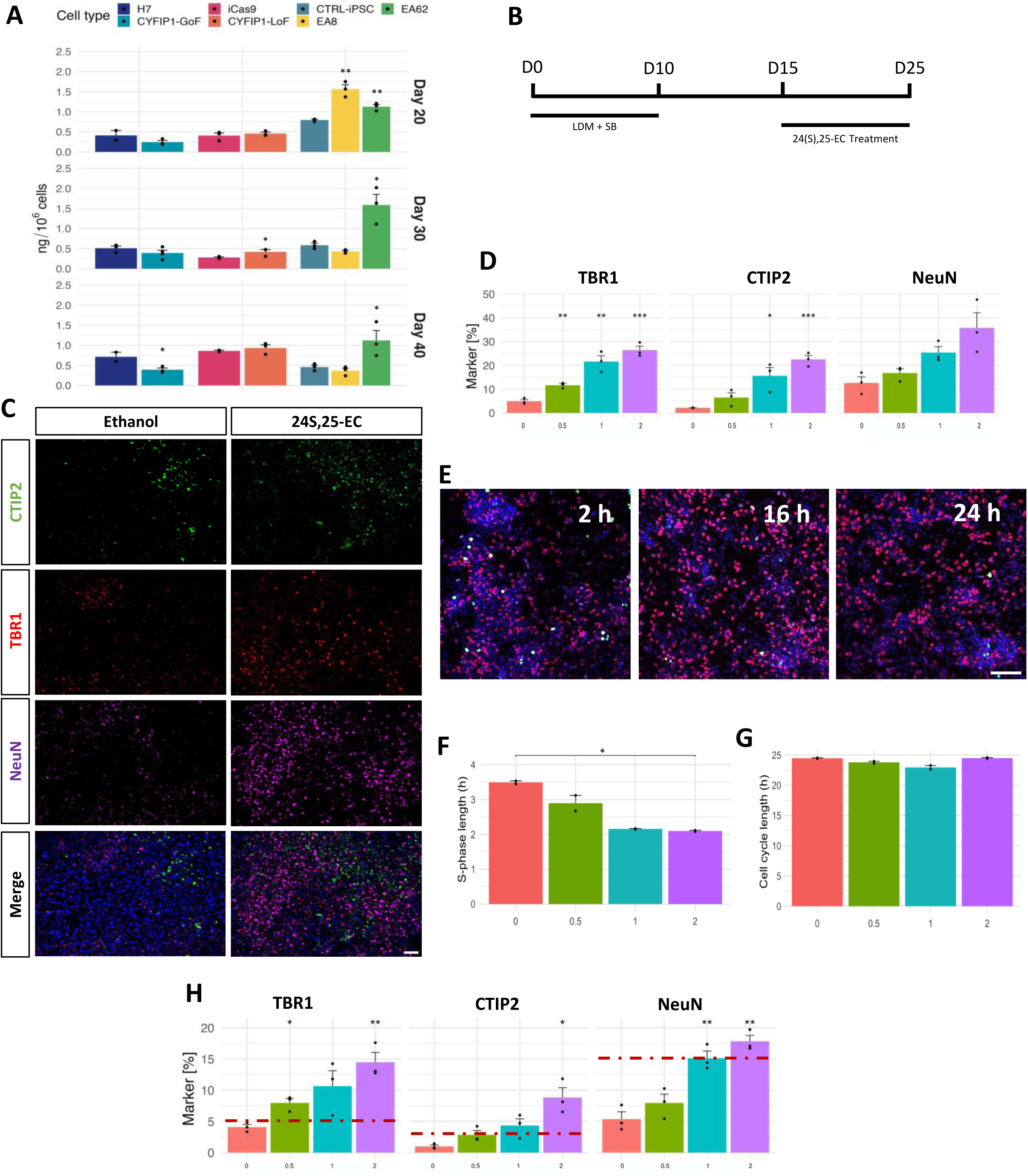
24S, 25-EC promotes neuronal differentiation of hESC-derived cortical NPCs. A) Bar graph showing the levels of 24S,25-EC produced by CYFIP1-GoF, CYFIP1-LoF, EA8 and EA62 15q11.2del iPSC and respective control PSC-derived neural cells at d20, 30 and 40. B) Schematic illustration showing 24S,25-EC treatment time windows. C) Representative images of immunostaining for TBR1, CTIP2 and NeuN at day 25 H7 cultures treated with ethanol vehicle control, and 24S,25-EC between day15-25. D) Quantification of the immunocytochemical analysis represented in C. E-G) Cumulative EdU incorporation assay of H7 cortical progenitor cultures treated with vehicle and increasing doses with 24S,25-EC from d15. Length of S-phase (F) and total cell cycle (G) determined by Nowakowski equation for all conditions. H) Quantification of TBR1^+^, CTIP2^+^ and NeuN^+^ in d25 CYFIP1-GoF cultures after 10 days exposure to increasing dose of 24S,25-EC or ethanol control. The red dashed line indicates the baseline (untreated) level in the isogenic control line. Data shown are mean±s.e.m. from three biological replicates per measurement (A) or treatment (D, F-H). Groups were compared by Student t-test between the indicated genotype and respective control per time point in A and one-way ANOVA followed by Tukey correction for D, F-H (*p<0.05; **p<0.01; ***p<0.001). Nuclei were counterstained with DAPI (blue). Scale bars: 50 µm.

### 24S,25-EC promotes neurogenesis and restores neuronal differentiation of CYFIP1-GoF cortical progenitors

24S,25-EC has been shown capable of promoting dopaminergic neuronal differentiation in hESC cultures and in the mouse brain^27–29^. Due to its increased levels in 15q11.2del and CYFIP1-LoF samples, we tested the hypothesis that 24S,25-EC can promote cortical neuronal differentiation which underlies the premature neurogenesis of 15q11.2del and CYFIP1-LoF NPCs. H7 hPSC-derived cortical cultures were exposed to 24S,25-EC for 10 days during d15-25 and d20-30, respectively, followed by immunostaining for TBR1, CTIP2 and NeuN (Figure 5B-D). Within the range of 0.5-2μM, we observed a dose-dependent increase of CTIP2^+^ neurons under both treatment regimes (Figure 5C-D, Figure S5A-B). 24S,25-EC exposure also elicited an increase of TBR1^+^ and NeuN^+^ neurons in both experimental setting although statistics significance was only reached under the d15-25 condition. This observation is consistent with the temporal order of TBR1^+^ and CTIP2^+^ neuron production during hPSC differentiation which is in line with the normal process of cortical neurogenesis. The neuronal promoting effect of 24S,25-EC was also observed in cortical neural cultures derived from iCas9 hESC line (Figure S5C-D).

We next performed a cumulative EdU incorporation assay to investigate whether 24S,25-EC enhances neuronal differentiation by regulating cell cycle kinetics of the responding cells. D20 H7 cortical cultures were exposed to 24S,25-EC or ethanol control from 2 to 48 hours followed by EdU staining. The cell cycle (Tc) and S-phase (Ts) length were then calculated using the Nowakowski equation^30^. 24S,25-EC treatment did not affect the Tc but resulted in a dose dependent decrease in Ts compared to ethanol controls (Figure 5E-G). Consistent with this finding, we observed a reduction of Ki67^+^ and Phospho-Histone H3^+^ (PH3^+^) cells after 3 days of 24S,25-EC treatment to d20 progenitors, indicating a decrease of total cycling cells (Figure S5E-F).

The neurogenic activity of 24S,25-EC observed above prompted us to ask whether this oxysterol can ameliorate reduced neuronal production of CYFIP1-GoF progenitors. We found that 0.5μM 24S,25-EC treatment of CYFIP1-GoF culture during d15-25 was able to restore the number of TBR1^+^ and CTIP2^+^ neurons to the baseline H7 control levels (Figure 5H). Higher doses of 24S,25-EC resulted in further increase of neuronal numbers in CYFIP1-GoF cultures. Interestingly, the fold changes of neuronal numbers to vehicle control in responding to increasing 24S,25-EC concentration were comparable between the CYFIP1-GoF and H7 isogenic control cultures despite the lower absolute neuron numbers in CYFIP1-GoF cultures in the basal condition or when treated with 24S,25-EC of the same doses (Figure 5H, Figure S5G)

### LXR signalling mediates oxysterol regulation of neurogenesis in hPSC-derived cortical progenitors

A number of oxysterols, including 24S,25-EC, have been identified as endogenous ligands of LXR^31, 32^, raising the question whether LXR signalling plays a role in CYFIP1-regulation of neurogenesis. LXRβ is the dominant isoform expressed in the brain and functions via forming obligatory heterodimer with retinoid X receptor (RXR)^33^. We therefore analyzed the RNAseq dataset further for differential expression of LXRβ, RXR and their target genes in 15q11.2del, CYFIP1-LoF and CYFIP1-GoF NPCs and neurons compared to their respective controls^34, 35^. We found that while no significant changes were detected in the transcript levels of LXRβ and RXR themselves, many of the LXRβ and RXR target genes were amongst the DEGs of CYFIP1-GoF, CYFIP1-LoF and the two 15q11.2del lines at both NPC and neuronal stages. Examples include master transcription factor for sterol biosynthesis (SREBF1), genes involved in neural progenitor proliferation and differentiation (ID1&2, HEY1, SMAD3&5) and G1 cell cycle phase progression (CDK4)(Table S1-2, Table S5-6). Fisher exact test confirmed significant enrichment of LXRβ target DEGs against the total non-LXR target DEGs in CYFIP1-GoF NPCs, CYFIP1-LoF NPC and neurons and EA8 iPSC NPCs, while RXR targets were also significantly enriched in CYFIP1-GoF NPCs and EA8 iPSC NPCs and neurons (Figure 6A). Thus, the transcriptomics data indicates altered LXR signalling in neural cells with 15q11.2del, CYFIP1-LoF and CYFIP1-GoF.

**Figure 6.**
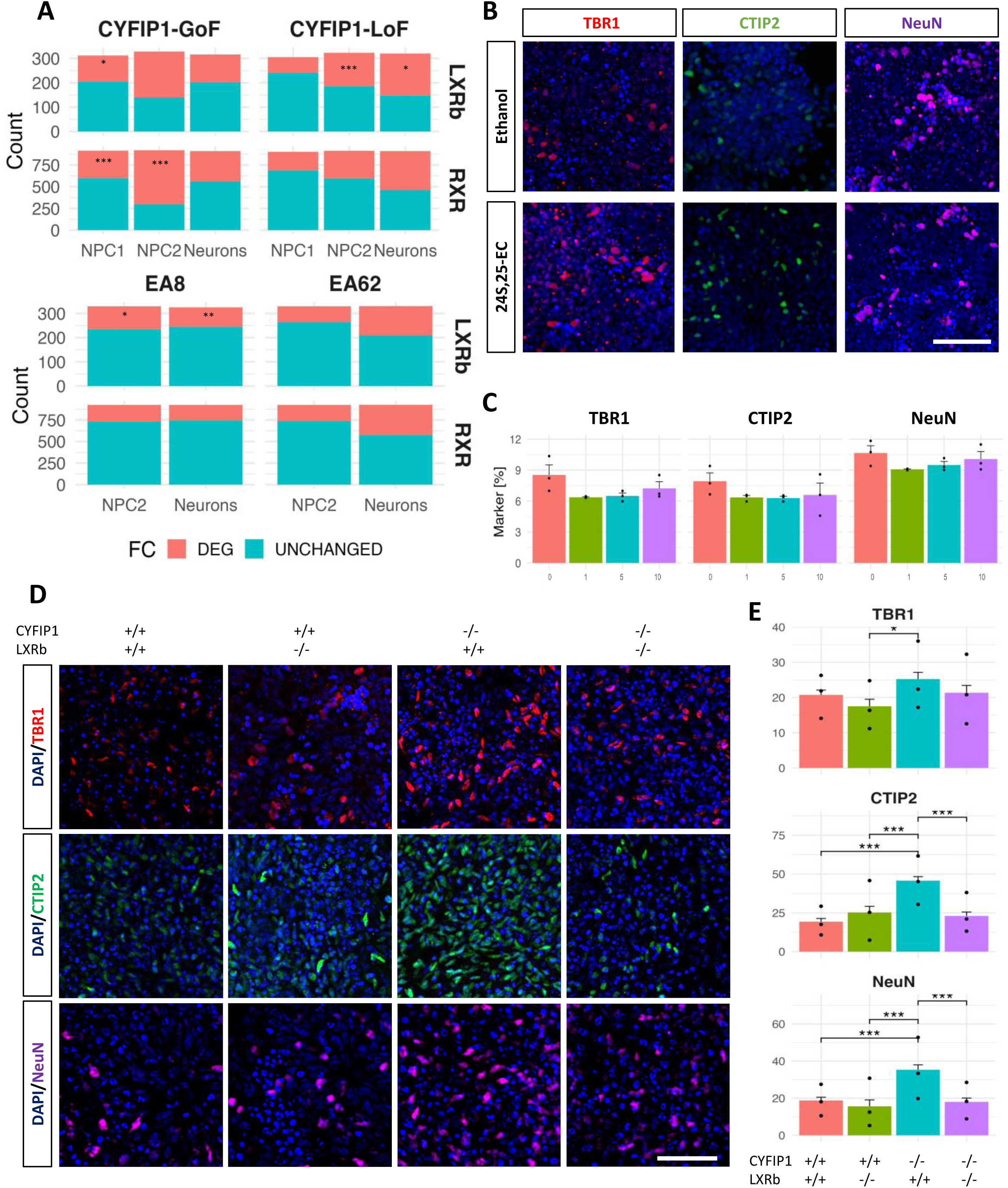
LXR signalling mediates oxysterol-regulated neurogenesis. A) Bar graph showing the number of differentially expressed LXRβ and RXR target genes in CYFIP1-GoF, CYFIP1-LoF and 15q11.2del NPCs and neurons. LXRβ and RXR targets were compared to non-target DEGs by one-sided Fisher’s exact test (*p<0.05; **p<0.01; ***p<0.001). B-C) 24S,25-EC treatment during d15-25 failed to promote neuronal production of LXRβ-deficient NPCs. D-E) Premature neuronal differentiation of CYFIP1-LoF NPC is reversed by compound deletion of LXRβ and CYFIP1. Data shown in C and E are mean±s.e.m. from three biological replicates per treatment (C) or genotype (E). Groups were compared by one-way ANOVA followed by Tukey correction (*p<0.05; **p<0.01; ***p<0.001). Nuclei were counterstained with DAPI (blue). Scale bars: 50 µm.

In order to provide direct evidence that LXR signalling mediates oxysterol regulation of neuronal differentiation, we generated a loss-of-function LXRβ (NR1H2) line via CRISPR/Cas9 genome editing in iCas9 hESCs (referred to as LXRβ-/-) and examined the response of LXRβ-/- NPCs to 24S,25-EC using the same experimental paradigm as described above (Figure 5, Figure S6). In contrast to that observed in H7 and iCas9 cultures treated with 24S,25-EC (Figure 5C-D, Figure S5), no differences in the number of TBR1^+^, CTIP2^+^ and NeuN^+^ neurons were detected between the vehicle control and 24S,25-EC treated LXRβ-/- cultures (Figure 6B-C). This finding demonstrates that LXRβ is required for 24S,25-EC regulation of neurogenesis.

### LXRβ knockout attenuates premature neuronal differentiation of CYFIP1-deficient cortical progenitors

24S,25-EC levels were increased in CYFIP1-LoF and 15q11.2del cells, accompanied by apparent over-activation of LXR signalling in these cells as suggested by the transcriptomics analysis. These findings point to the hypothesis that dysregulated LXR signalling underlies the premature neuronal differentiation and its blockade would rescue or partially rescue this deficit. To this end we also genetically deleted LXRβ in a CYFIP1-LoF line (CYFIP1-/-LXRβ-/-) and examined the consequence of CYFIP1/LXRβ compound deletion on neuronal production in comparison with the isogenic trios of CYFIP1-/-, LXRβ-/- and iCas9 hESCs.

We found that LXRβ single knockout didn’t affect the number of TBR1^+^, CTIP2^+^ and NeuN^+^ neurons compared to its isogenic iCas9 control (Figure 6D-E). As expected, neuronal production was increased in CYFIP1-LoF cultures at d30. However, the double knockout cultures contained a comparable number of neurons to that of the iCas9 cultures (Figure 6D- E), indicating that ablation of LXRβ in CYFIP1-/- neural progenitors attenuated the premature neuronal differentiation due to CYFIP1 deficiency. This result demonstrates that LXR signalling serves as a mechanism underlying the aberrant neurogenesis in CYFIP1 and 15q11.2del cortical cultures.

## Discussion

The notion that changes in temporal dynamics of developmental processes underlie neurodevelopmental disorders is gaining momentum^36–39^, but precise molecular mechanisms have remained largely unknown. This study identifies CYFIP1 regulation of cholesterol biosynthesis as a novel regulatory axis playing an important role in neurogenesis, and suggests that dysregulation of LXR signalling consequent of 15q11.2CNV may contribute to the pathogenesis of associated neurological disorders.

### CYFIP1, a new regulator of neurogenesis and causal risk gene in 15q11.2 loci

Being part of the WAVE regulatory complex and binding partner of FMRP, respectively, CYFIP1 plays an important role in actin cytoskeleton formation and protein translation. Accordingly, CYFIP1 functions have been reported in neural progenitor migration^20^, brain structure^18, 19^ and dendritic morphology, synaptic activity and associated behaviors^10, 12–15, 40^. Recent mouse genetic studies also revealed a role for Cyfip1 in adult neurogenesis although opposite effects were reported^16, 17^. The underlying molecular mechanisms by which Cyfip1 affects adult neural stem cells in these mouse models remain unknown. The current study presents the first demonstration that CYFIP1 is required for neurogenesis early on in the embryonic stages, a cellular function in line with its expression during human cortical development. Both CYFIP1-deficiency and its elevated expression led to disturbed neurogenesis, suggesting that a tight control of CYFIP1 dose is crucial for coordinated neural progenitor proliferation and terminal differentiation. Inappropriate neuronal numbers and/or birth time of a defined subtype may result in errors in their final destinations and mismatched neuronal interconnections in the brain, which may contribute to the mirror phenotypes in MRI studies of white matter microstructure and the network dysfunction present in patients with neurodevelopmental disorders attributed to CNVs at the 15q11.2 locus^41^.

CYFIP1 is generally considered as the major risk gene within the 15q11.2 locus that impacts on adverse neurological outcomes^3, 5, 6, 42^, but direct evidence for this has been lacking. Using the same neuronal differentiation paradigm, we show that NPCs derived from 15q11.2del iPSCs present a closely analogous phenotype to hESC-derived NPCs deficient in CYFIP1 alone. Consistent with the experimental findings, genome-wide transcriptomic analysis revealed significant dysregulation of transcripts related to neurogenesis from both CYFIP1-modified hESC- and 15q11.2del iPSC-neural derivatives, and crucially a high degree of overlap in the biological processes (GO terms), despite the variations between independent carrier iPSC lines and that CYFIP1-GoF and CYFIP1-LoF cells were derived in different parental lines. Therefore, our work provides a direct demonstration for CYFIP1 as the likely causal risk factor of 15q11.2 deletion effects on neurogenesis.

### Oxysterol regulation of cortical neurogenesis and implications in pathogenesis of neurodevelopmental disorders

The works presented here identified a pro-neuron differentiation activity of 24S,25-EC in human cortical progenitors. We have shown previously that 24S,25-EC is abundantly produced in the developing mouse cortex^43^, while Wong et al detected this oxysterol also in primary cultured human fetal cortical neurons^44^. We found in this study that 24S,25-EC is already produced by day 20 human NPCs at comparable levels to that of day 30-40 neuronal cultures. These findings together support a physiological role of 24S,25-EC in regulating cortical NPC differentiation in the fetal brain, as it has been shown previously in the developing ventral midbrain^28, 29^.

Does 24S,25-EC target NPCs promiscuously or defined progenitor subtypes? In the embryonic cortex, neurons are produced from both apical and basal progenitors while apical progenitors also give rise to basal progenitors. The human brain generates more basal progenitors during development as a strategy to producing more neurons. Basal progenitors have a longer cell cycle (Tc) than apical progenitors while the expanding (proliferating) neural progenitors, both apical and basal, shorten S-phase length (Ts) when they commit to neuronal production^45^. 24S,25-EC shortens the Ts of hESC-derived cortical progenitors without affecting Tc, suggesting it may promote the transition from expanding neural progenitors to neurogenic progenitors, regardless of they are apical or basal progenitors.

The discovery that 15q11.2del and CYFIP1 dosage change led to abnormal 24S,25-EC formation presents the first causal link between autism and schizophrenia risk CNV and distorted sterol biosynthesis, while the ability of 24S,25-EC and genetic blockade of its downstream signaling (via LXRβ knockout, see discussion below) in restoring delayed and premature neuronal differentiation of CYFIP1-GoF and CYFIP1-LoF NPCs, respectively, implicates a potential role for distorted sterol biosynthesis in the pathogenesis of 15q11.2CNV associated disorder. It would be interesting to investigate whether sterol dysregulation also plays a role in neurodevelopmental disorders associated with other risk genes/loci. Relevant to this hypothesis, changes in SREBP2 and CYP46A1 protein levels have been reported in several brain regions in a preclinical model of Fragile X Syndrome^46^.

A role for oxysterols in pathogenesis of neurodevelopmental disorder was also suggested by a recent study of mouse and hESC models of Smith-Lemli-Opitz syndrome (SLOS)^47^. SLOS is caused by mutations in the cholesterol biosynthesis gene 7-dehydrocholesterol reductase (DHCR7) that converts 7-dehydrocholesterol (7-DHC) to cholesterol. SLOS patients present a host of neurodevelopmental abnormalities including small head size (microcephaly), severe developmental delay and features of autism^48^. Loss of Dhcr7/DHCR7 alleles results in decreased proliferation and increased neurogenesis in the developing mouse cortex and hESC-derived cortical progenitors, caused by an accumulation of 7-dehydrocholesterol (7-DHC) and 7-DHC-derived oxysterols^47^.

Several recent studies highlighted the potential for brain-derived oxysterols as biomarkers of neurodegenerative conditions such as Alzheimer’s disease^49^. Unlike cholesterol, oxysterols readily traverse the blood brain barrier from the brain to periphery and vice versa, thus plasma levels of brain-derived oxysterols reflect cholesterol biosynthesis and metabolism in the brain^50^. Altered oxysterol levels have been described in patients with several neurodegenerative disorders, including Alzheimer’s disease, Amyotrophic Lateral Sclerosis, Parkinson’s disease, and Huntington’s disease^51–53^. While investigations into sterol dysregulation in neuropsychiatric and neurodevelopmental disorders is in its infancy, changes of plasma 24S-hydroxycholesterol, a primarily neuron derived oxysterol, is found in patients with schizophrenia^54^, ASD^55^, and bipolar disorders^56^.

### Molecular mechanisms underlying CYFIP1 regulation of neurogenesis

Accelerated or delayed neurogenic differentiation, often as exemplified by alterations of neural progenitor to neuron ratio during early neurogenesis, have been reported in multiple studies of human stem cell models carrying neurodevelopmental risk variants^36–39, 57^. However, the molecular mechanisms responsible for the disrupted process remain largely unknown. By demonstrating the loss of pro-neuron activity of 24S,25-EC and the rescue of premature neuronal differentiation of CYFIP1-LoF NPCs by genetic deletion of LXRβ, respectively, the works described here establishes LXR as the overriding signalling mechanism underlying the disturbed neurogenesis in CYFIP1-LoF. This finding is consistent with previous report that 24S,25-EC is a potent endogenous ligand of LXR as well as being the most abundant LXR ligand over other oxysterols in the developing brain playing a role in midbrain dopaminergic neurogenesis in vivo^27, 29, 43^.

Several lines of evidence support a direct regulatory role of CYFIP1 in cholesterol biosynthesis. CYFIP1 is known to regulate protein translation via binding to mRNA targets as part of the protein complex with FMRP and elF4E^10, 58^. SREBP1 and SREBP2 (Sterol Regulatory Element Binding Transcription Factor 1 & 2, respectively) are master transcription factors for genes regulating lipid homeostasis. SREBP1 regulates lipogenic genes, whereas SREBP2 preferentially activates genes of cholesterol biosynthesis and metabolism^59, 60^. SREBP2 RNA has been shown recently a direct target of CYFIP1 in the adult mouse cortical and hippocampus tissues via RNA immunoprecipitation sequencing (RIP-seq) against CYFIP1^40^. Moreover, SREBP2 was also identified as a high-confidence FMRP target in the brain, together with RNAs for two enzymes in the cholesterol synthesis pathway: HMGCS1 (Hydroxymethylglutaryl-CoA synthase) and GGPS1 (geranylgeranyl diphosphate)^61, 62^ while SREBF1 and 14 other genes in the cholesterol synthesis pathway are reported to bind to FMRP in HEK cells^63^. In line with the above, majority of genes involved in cholesterol biosynthesis and metabolism are differentially expressed, mostly in opposite direction of change in CYFIP1-GoF and LoF neural progenitors and neurons (Figure 3F) and the transcriptomic changes are in line with the alteration of 24S,25-EC in neural cells of respective genotypes (Figure 5A). Together these findings provide mechanistic link between CYFIP1 dosage change and alteration of 24S,25-EC profile in 15q11.2del and CYFIP1 disrupted PSC neural derivatives.

In summary, we report a new role for CYFIP1 in the 15q11.2 CNV in human neurogenesis, which is mediated by oxysterol-LXR signalling axis through alterations in cholesterol biosynthesis. This novel mechanistic insight opens fresh perspectives to approach neuropsychiatric and neurodevelopmental disorders. Given the potential of oxysterols as disease biomarkers and druggable targets, this study may shed light into the development of novel diagnostic and therapeutic avenues in associated neurological disorders.

## Materials and Methods

### Stem cell culture and differentiation

Human pluripotent stem cells (hPSCs) lines used in this study include H7 (WiCell), iCas9 that conditionally express Cas9 upon doxycycline stimulation^22^, and iPSCs derived from two male 15q11.2 deletion carriers and two healthy control individuals. CYFIP1-GoF cells were established by integrating a human CYFIP1 expression cassette on H7 background while CYFIP1-LoF cells were generated by CRISPR/Cas9 assisted gene targeting using iCas9 as the parental line. Details on the generation of these edited lines are provided in supplementary information. All hPSCs were cultured on matrigel coated plastics in E8 media (ThermoFisher). Media was changed daily and cells passaged mechanically with 0.02% EDTA at 80% confluence. Cortical neuronal differentiation was performed as described previously^57, 64^. Briefly, PSCs from 80% confluent wells of a 6-well plate were plated onto a 12-well plate previously coated with reduced growth factor matrigel (VWR) in E8 media (d0) and changed to retinoid free N2B27 the next day. For the first 10 days, cultures were supplemented with SB431542 (10µM, Tocris) and LDN-193189 (100nM, Tocris). During differentiation the cultures were split twice using EDTA. The first split was done *en bloc* onto fibronectin-coated plates at approximately d10 and the second split to single cells onto poly-D-lysine/laminin-coated coverslips at around d20. Retinol free B27 was switched to standard B27 on d20 to aid neuronal maturation. For oxysterol experiments, 24S, 25-epoxycholesterol (1-10μM, Abcam) or ethanol control were added to N2B27 media for 10 days either during d15-25 or d20-30.

### Immunocytochemistry

Cultures were fixed with 3.7% PFA for 15-20 min at 4 °C. For nuclear antigen detection an additional fixation with methanol gradient was performed, which include 5 mins each in 33% and 66% methanol at room temperature followed by 100% methanol for 20 min at -20. Cultures were then returned to PBST via inverse gradient and were then permeabilized with three 10-minute washes in 0.3% Triton-X-100 solution in PBS (PBS-T) and then blocked in PBS-T containing 1% BSA and 3% donkey serum. Cells were incubated with primary antibodies in blocking solution overnight at 4°C. Following three PBS-T washes, Alexa-Fluor secondary antibodies (Thermo Fisher Scientific) were added at 1:1000 PBS-T for 1 hour at ambient temperature in the dark. Three PBS-T washes were then performed that included once with DAPI at 1:1000 (Thermo Fisher Scientific). Images were taken on a Leica DMI6000B inverted microscope or a PerkinElmer Opera Phenix. Quantification was carried out in Cell Profiler (cellprofiler.org) or manually using ImageJ (imagej.net) by examining randomly selected fields from at least 3 independent experiments. The antibodies used are provided in the supplementary table 9.

### EdU Labelling and detection

Cells in S-phase were labelled using the Click-iT EdU Assay kit (Thermo Fisher). For EdU pulse analysis, cells were incubated with EdU for 2 hours at specified day of differentiation. For EdU cumulative assay required for calculating cell cycle length, cells were incubated for increasing periods of time of 2,4, 8, 16, 24 and 48 hours. Cells were then fixed in 3.7% PFA for 15 min at 4 °C. EdU detection was carried out as per manufacturer’s protocol. Cell cycle length determination was performed using the Nowakowski equation^30^. Nine fields each from 3 biological replicates were counted at each time point of the EdU cumulative labelling. The percentages of EdU^+^ in ethanol and 24S,25-EC conditions were plotted against time. A linear regression was performed, and the values of the slope (m) and y-intercept (b) of the linear equation were used to calculate the values of cell cycle length (tc) and S-phase length (ts), using the Nowakowski equation:

GF(t) = (GF/tc)t + GF(ts/tc). Where: GF= Growth fraction in cell population, m=GF/tc thus tc=GF/m, b= GF(ts/tc) thus ts=(b*tc)/GF

### Flow cytometry

Cultured cells were dissociated in accutase (ThermoFisher) for 10 minutes at 37 °C, then washed and counted. Samples containing 10^6 cells were fixed in 70% EtOH for 2 hr at 4 °C. The cells were washed in DPBS and incubated with DAPI (0.3 µg/ml, ThermoFisher) and of RNaseA (200 µg/ml, ThermoFisher) for 30 minutes. The samples were analysed on a BD LSRFortessa cell analyser (BD Biosciences). Results were analysed using FLOWJO software (Tristar, Ashland, OR, USA). Statistical analysis was performed in R (www.r-project.org).

### Analysis of RNA sequencing data

Details on RNA extraction and RNA sequencing are provided in Supplemental Information. FASTQ files were trimmed and mapped to the Ensembl human genome GRCh38.84 (hg38) using STAR 2.5.1b.65. Quality of the samples was assessed using FastQC (v 0.11.2) prior and after trimming. Gene counts were obtained using the Samtools Feature count command. Library size normalisation and differential gene expression was carried out using the R/Bioconductor package DESeq2. Genes with less than 10 counts in at least one of the replicates were excluded for the analysis. Subsequent study of the DEGs was performed on protein-coding Entrez genes with a Benjamini Hochberg-corrected p<0.05. The R/Bioconductor package clusterProfiler66 was used to identify GO terms enriched in DEGs when compared to a background set consisting of all protein-coding genes expressed in the cell-line at that developmental stage. To remove highly similar GO terms, semantic similarity between terms was calculated using the Wang method. Terms with a threshold over 0.7 were removed keeping the term with the smallest p value.

### Western Blot

Cultured cells were lysed on ice using RIPA buffer (Abcam) supplemented with protease and phosphatase inhibitors (Sigma). Cell lysates were centrifuged for 15 minutes at 12000g and the resulting supernatant was combined with 1X Bolt LDS Sample Buffer (ThermoFisher) and 1X Bolt Sample Reducing Agent (ThermoFisher) and boiled at 97°C for 5 minutes. Equal amounts of proteins for each sample were separated on 4-12% Bolt Bis-Tris Plus gels (ThermoFisher) and then transferred to a PVDF membrane (0.45 µm pore size, Amersham Hybond, GE Healthcare) via electro-blotting. The membrane was blocked in 5% BSA (Sigma) and incubated with primary antibodies overnight at 4°C. The membrane was then washed in TBS-T (Tris Buffered Saline containing 0.1% Tween). Secondary antibodies were incubated for 2 hours at room temperature. The membranes were imaged in an Odyssey CLx Imaging system (LI-COR) or in a Gel Doc XR system (Bio-Rad). A list of the primary antibodies used for this study can be found in supplementary table 10.

### Sterol extraction and LC-MS analysis

Sterols and oxysterols are extracted from cell pellet using ethanol containing deuterated internal standards. The extracted oxysterol and sterol fractions were derivatized with Girard P reagent and analyzed by LC-MS(MSn) on a Orbitrap ID-X mass spectrometer. Mass spectra were recorded at high resolution in the Orbitrap analyzer, and MSn spectra recorded simultaneously in the ion-trap. Quantification was performed by stable isotope dilution on reconstructed ion-chromatograms generated in the Orbitrap. Details for the extraction and internal standards are provided in Supplemental information.

### Statistical analyses

All data were collected from at least three independent experiments and presented as mean± s.e.m. unless otherwise specified. Data were tested for normality with the Shapiro Wilk test and for equal variance with Levene test before performing statistical analyses by *t*-test or ANOVA as stated in the figure legends where relevant. Post-hoc Tukey test was applied following ANOVA for multiple comparisons. All statistical tests were performed in R (www.r-project.org).

**Additional methods and gene expression data are provided in supplemental information (link upon publication)**

## Acknowledgements

We thank members of the M.L. laboratory and the DEFINE community for helpful discussions during the course of this study. Thanks also to Dr. Craig Joyce for his contribution in the generation of 15q11.2 iPSC lines and Dr. Joanne Morgan for the support on RNA sequencing. RNA sequencing analysis was performed using the computational facilities of the Advanced Research Computing@Cardiff (ARCCA) Division, Cardiff University. The data and samples of the DEFINE cohort were collected and curated by Dr Stefanie Linden, Dr Faraz Ali, Kali Barawi, Ffion Evans, Jacqueline Smith, Alister Baird and Rachael Adams, with support from National Centre for Mental Health. This work was supported by a Wellcome Trust Strategic Award (100202/Z/12/Z) to M.O., J.H., D.L., M.L. and other co-PIs and fundings from the Medical Research Council (MRC, grant no. MR/R022429/1 to M.L.). Work in Swansea was supported by funding from the Biological Sciences Research Council (BBSRC, grant nos, BB/S019588/1 to W.J.G., BB/L001942/1 to Y.W.), the European Union, through European Structural Funds (ESF), as part of the Welsh Government funded Academic Expertise for Business project (to W.J.G. and Y.W.) and the Wellcome Trust Strategic Award (100202/Z/12/Z) through M.L. E.S. is funded by the BBSRC SWBio DTP. For the purpose of open access, the author has applied a Creative Commons Attribution (CC BY) licence to any Author Accepted Manuscript version arising from this submission.

## Author Contribution

D.C. F., C.T. and M.L. conceived the study and designed the experiments. D.C. F. and C.T. carried out and analyzed the hESC and hiPSC experiments. D.C.F. performed RNAseq data analysis with support from A.P. W.J.G and Y. Wang conceived the sterol profiling study while E.S. performed the experiments and analyzed the data. D.L., J.H. and M.O. provided 15q11.2del carrier samples and contributed to general discussions throughout the work. M.L. C.T. D.C.F. wrote the paper. All authors edited and approved the paper.

## Availability of data and materials

- The raw RNA sequencing data generated during the current study is available in the GEO repository, accession number (to be provided here in publication).
- The datasets used and/or analyzed during the current study are available from the GitHub repository (link to be provided here in publication) and the corresponding author on reasonable request.
- All data generated or analyzed during this study are included in this published article (and associated supplementary information files).

## Supplementary figure legends

**Figure S1.**
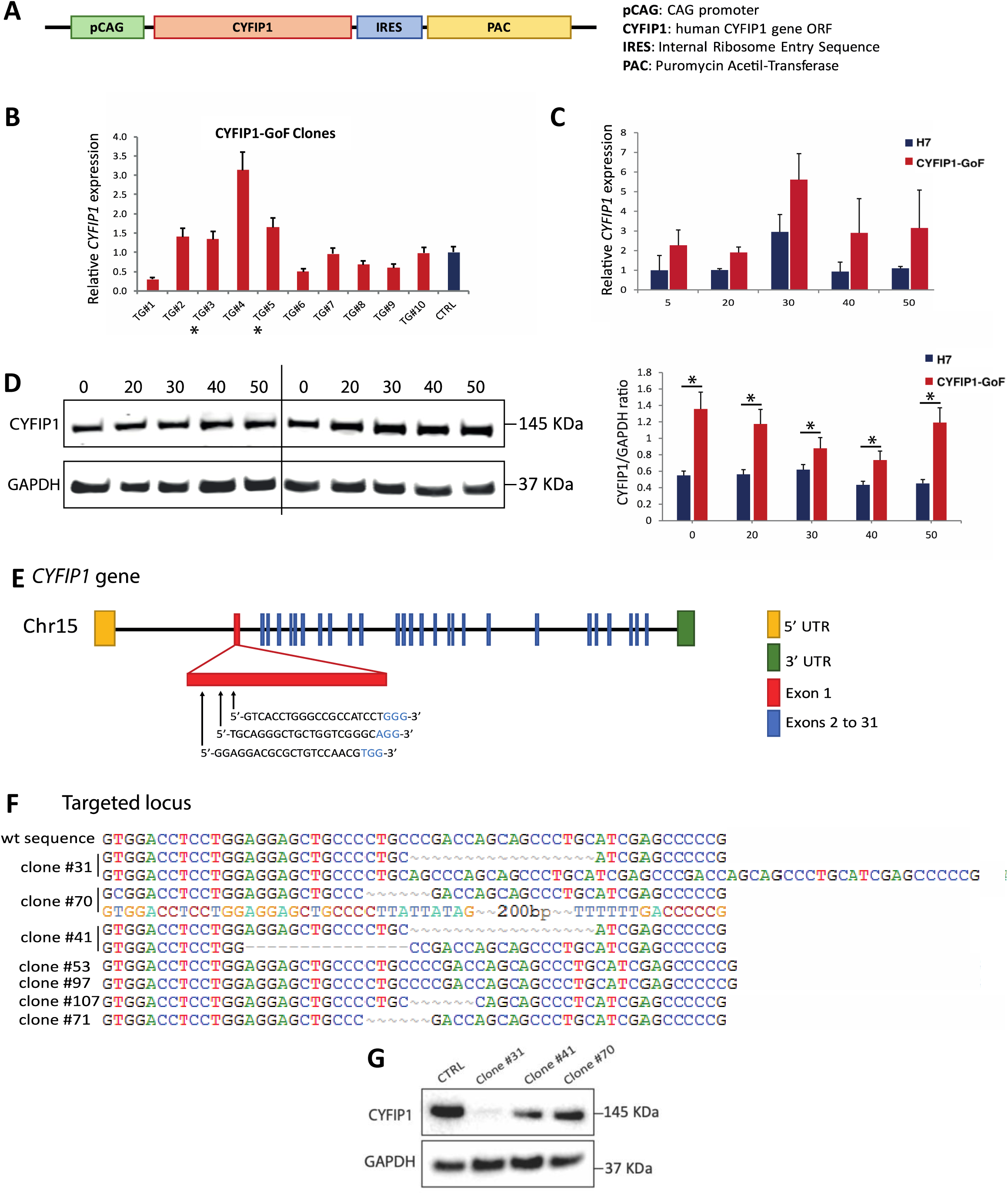
Derivation of CYFIP1-GoF and CYFIP1-LoF hESCs lines. A) Schematic representation of human CYFIP1 expression vector. B) qPCR quantification of CYFIP1 transcript levels in independent transgenic CYFIP1-GoF hESCs lines (TG#1 -10) and parental H7 cells (CTRL) at undifferentiated state. Data were normalised to the CTRL which was set as 1. C) qPCR analysis of CYFIP1 transcript during neuronal differentiation. Data shown are the mean between clones #3 and #5, all normalised to the CTRL cells at d5 of neuronal differentiation. D) Western blot confirming increased CYFIP1 protein level in CYFIP1-GoF cells (Clone #3) throughout 50 days of neuronal differentiation. Quantitative data on the right panel represent the average of clones #3 and #5. E) Schematic of *CYFIP1* gene locus highlighting the first exon where gRNA sequences were designed. For space reasons, the sequence of the insertion present in one of the alleles of clone#70 is not reported and is indicated as “--200bp--”. F) DNA sequence flanking the edited site in the wild-type and edited CYFIP1 locus. G) Western blot demonstrating reduced CYFIP1 protein levels in 3 of the targeted clones (CYFIP1 clone#31, clone#41 and #70) compared to the parental line (CTRL).

**Figure S2.**
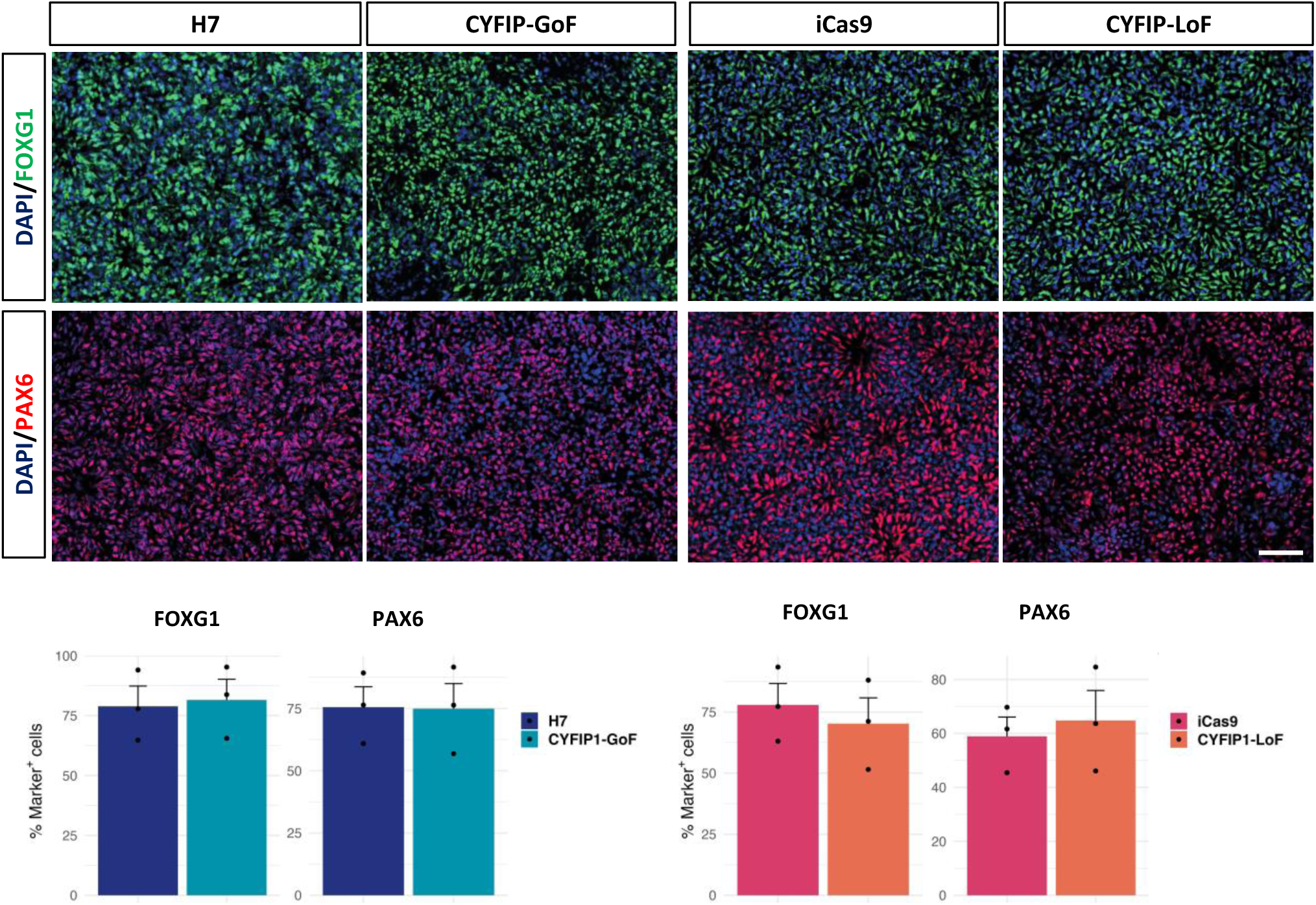
Cortical fate induction is not affected by gain- or loss-of CYFIP1. A-B) Immunostaining for PAX6 (red) and FOXG1 (green) in d15 CYFIP1-GoF and CYFIP1-LoF NPCs cultures and respective controls. C-D) Quantification of PAX6^+^ and FOXG1^+^ cells. Data represent mean±s.e.m. from three biological replicates and analysed by Student t-test. No significant differences were found. Nuclei were counterstained with DAPI (blue). Scale bars: 50 µm.

**Figure S3.**
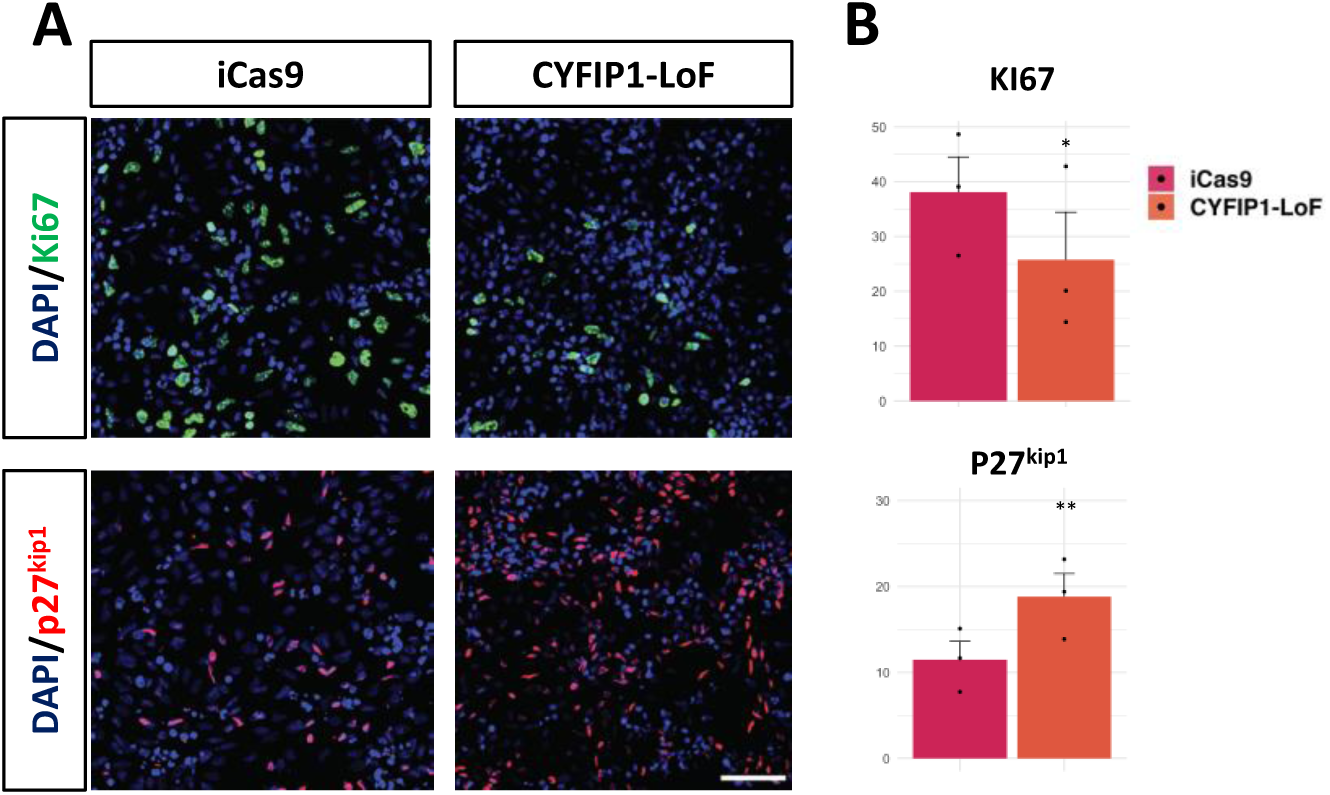
Premature cell cycle exit of CYFIP1-LoF progenitor cells. Immunostaining and quantification for the proliferation marker Ki67 (green) and the cell cycle regulator p27^kip1^ (red) in CYFIP1-LoF and isogenic iCas9 control progenitor populations. Nuclei were counterstained with DAPI (blue). Scale bars: 50 µm. Groups were compared by Student t-test (*p<0.05; **p<0.01).

**Figure S4.**
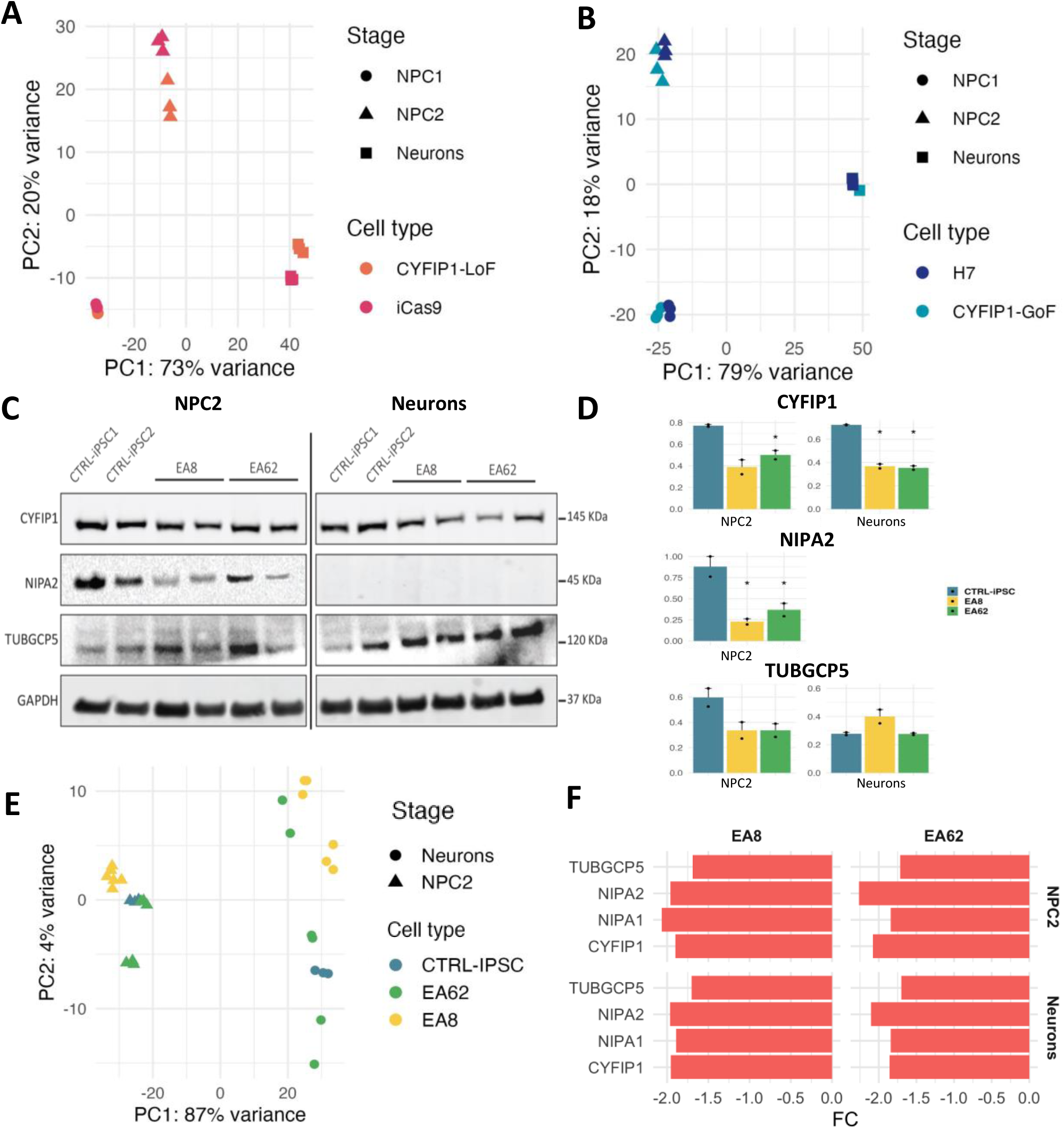
Additional information on RNAseq analysis and protein level of 15q11.2 genes. A-B) PCA plot of CYFIP1-LoF, CYFIP1-GoF and their respective isogenic control cultures at d10 (NPC1), d20 (NPC2) and d40 (Neurons). C-D) Western blots analysis for CYFIP1, NIPA2, TUBGCP5 and GAPDH (internal control) on neural cells derived from control (CTRL) and 15q11.2-deleted iPSCs (EA8 and EA62), at the NPCs and neuronal stage. The quantitative protein level is expressed as intensity of the band corresponding to the protein of interest, normalised to the intensity of the GAPDH band on the same gel. Groups were compared by one-way ANOVA, followed by Tukey post-hoc test (*p<0.05). E) PCA for NPCs and neurons derived from 15q11.2del iPSC (EA8 and EA62) and two control iPSC lines. PC1 captures the largest difference segregating the different developmental stages while PC2 captures differences between genetic backgrounds. C) Bar graphs showing relative levels of 15q11.2 CNV transcripts to respective control cells at each of differentiation stage analysed. Bar height represents fold change (FC)

**Figure S5.**
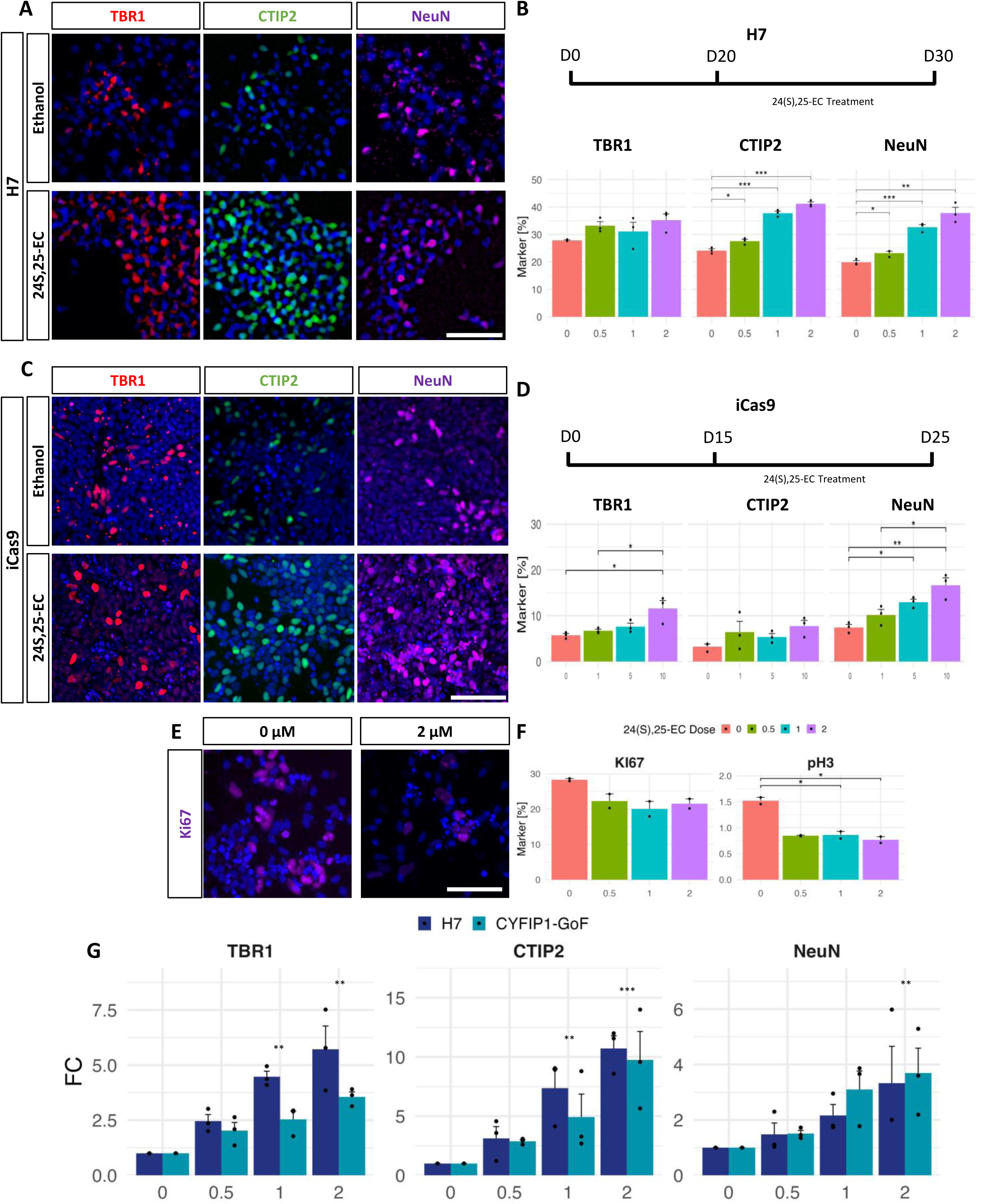
24S, 25-EC promotes neuronal differentiation of hESC-derived cortical NPCs (additional data) A-B) Illustration of experimental scheme, immunocytochemistry and quantification for TBR1, CTIP2 and NeuN in d30 H7 cultures treated with 24S,25-EC from d20. C-D) Illustration of 24S,25-EC treatment scheme for iCas9 iPSC line, immunostaining and quantification for TBR1, CTIP2 and NeuN in d25 iCas9 cultures. E) immunocytochemistry for Ki67 4 days after 24S,25-EC or ethanol exposure. F) Number of Ki67^+^ and phosphorylated histone H3^+^ (pH3^+^) cells in H7 cultures treated with or without increasing dose of 24S,25-EC for 4 days from d15. G) Quantification of TBR1^+^, CTIP2^+^ and NeuN^+^ in d25 CYFIP1-GoF and H7 control cultures after 10 days exposure to increasing dose of 24S,25-EC or ethanol control. Data shown are mean±s.e.m. from three biological replicates. Shown in ‘G’ are the same data of Fig 5H presented as fold change of 24S,25-EC treated cultures to none treated cultures of CYFIP1-GoF (light blue) and isogenic H7 control (dark blue), respectively. Groups were compared by one-way ANOVA followed by Tukey correction (*p<0.05; **p<0.01; ***p<0.001). Nuclei were counterstained with DAPI (blue). Scale bars: 50 µm.

**Figure S6.**
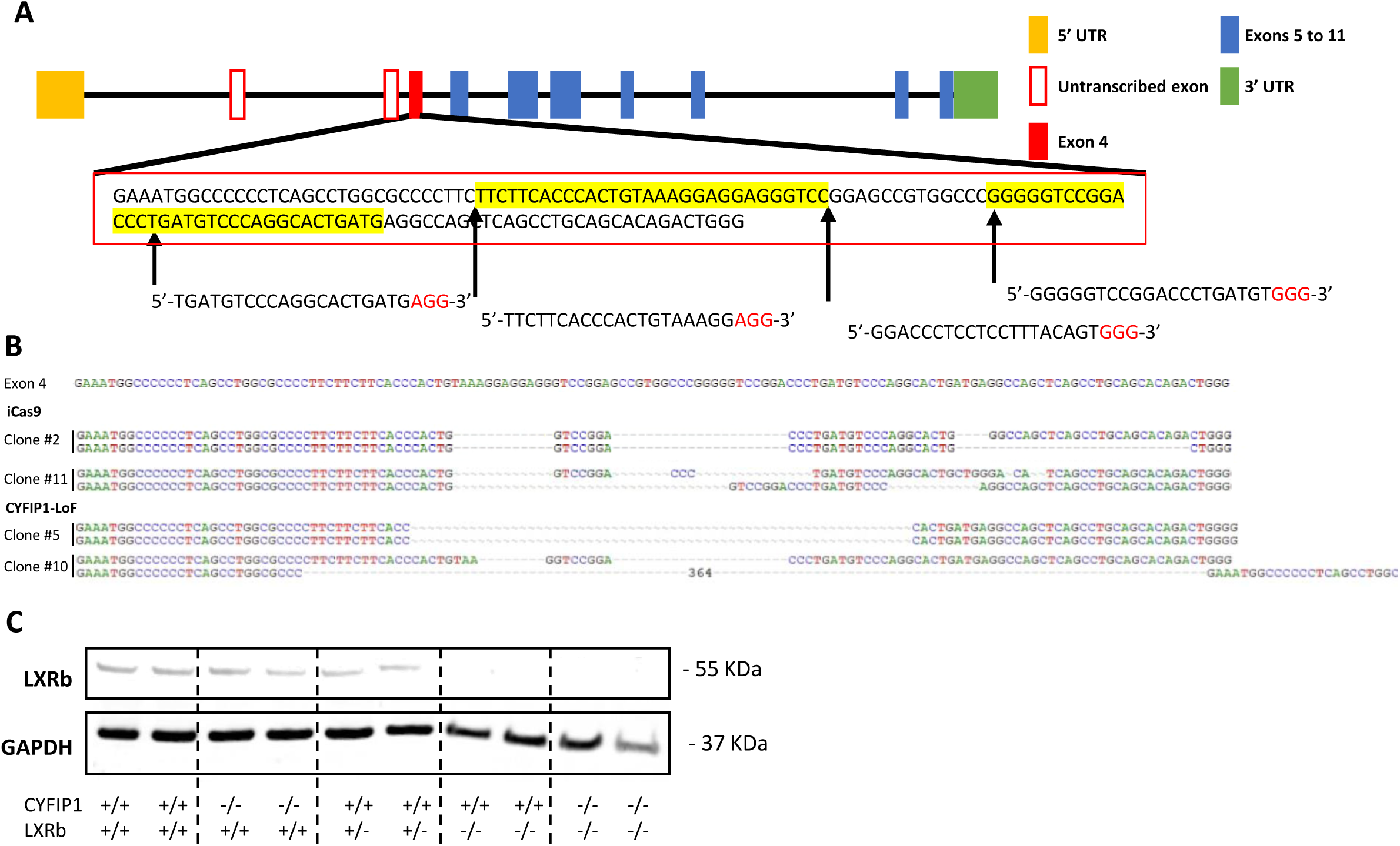
Generation of LXRβ knockout hESC lines in iCas9 and CYFIP1-/- hESCs. A) Schematic of LXRβ gene locus highlighting exon 4 where gRNA sequences were designed. B) Detection of InDels (insertions and deletions) via Sanger sequencing of the targeted exon 4 in iCas9 and CYFIP1-LoF background, respectively. C) Western blot analysis for LXRβ protein in iCas9 isogenic control, CYFIP1-LoF, LXRβ knockout and CYFIP1 & LXRβ double knockout iPSCs.

## Supplementary Materials and Methods

### Generation of CYFIP1-GoF and CYFIP1-LoF cells by transgene expression and CRISPR/cas9 assisted genome editing

For CYFIP1 overexpression (CYFIP1-GoF), human CYFIP1 open reading frame (SC100426, OriGene) was cloned into a pCAG-IRES-PAC plasmid upstream of the IRES sequence [1]. The resulting pCAG-CYFIP1-IRES-PAC vector was transfected into H7 hESCs using the 4D Nucleofector system (Lonza). Stable H7 transfectants were selected by Puromycin (1µg/mL, Sigma) for 10 days prior to colony picking and clonal expansion. CYFIP1 expression was validated by RT-PCR and Western blot analysis (figure S1). Two clones (#3 and #5) that exhibit approximately two-fold CYFIP1 protein level to that of the parental control H7 cells were used for this study.

For generating CYFIP1-LoF cells, guide RNAs (gRNAs) were designed to target the first coding exon, common to 6 of 7 isoforms of the *CYFIP1* gene (gRNA1 5’-GGAGGACGCGCTGTCCAACG -3’, gRNA2 5′-TGCAGGGCTGCTGGTCGGGC-3’, gRNA2 5′-GTCACCTGGGCCGCCATCCT-3’). All gRNAs were synthesized as RNAs by *in vitro* transcription and transfected into iCas9 hESCs as described by Gonzalez *et al* [2]. 110 individual colonies were picked from day 10 post transfection and clonally expanded while the remaining pool of transfected cells were used for Surveyor Assay (Integrated DNA Technologies). Genomic DNA was collected for each clone and the targeted region was screened by PCR of genomic DNA using two pairs of primers: Forward 1, 5’-ATGTGTTGTTCCAGCCCAGG-3’, Reverse 1 5’-CATCATGTGGGGTCGGAGC-3’, which give rise to a predicted 190bp product. Forward 2 5’-CCCTGAGAGAGACACGCAAC-3’, Reverse 1 5’-CCTCACTGCATAGTCTATTGGGA-3’, which give rise to a 557bp product. PCR products suggesting deletions and insertions (INDELs) were then confirmed by cloning into a pGEM-T Easy vector (Promega) followed by Sanger sequencing (GATC Biotech).

### Quantitative PCR

Total RNA was extracted using TRI reagent (Sigma) according to the manufacturer’s instructions. Reverse transcription was performed using SS RTIII (Invitrogen). QPCR was carried out using SYBR MESA green on a BioRad CFX connect system. All data were normalised to three reference genes (*GAPDH, b-ACTIN, CYCLOPHILIN*). The sequence information for PCR primers are provided below. All qPCR data are presented as mean ±s.e.m. of biological duplicates or triplicates.

#### Primers used for qPCR

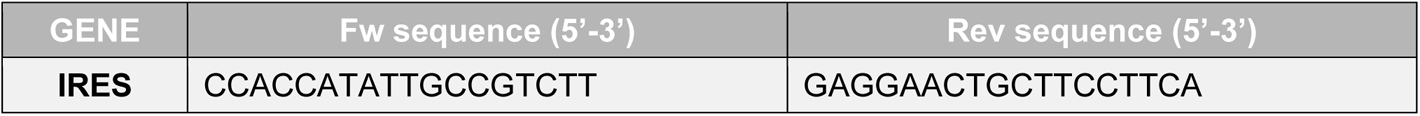
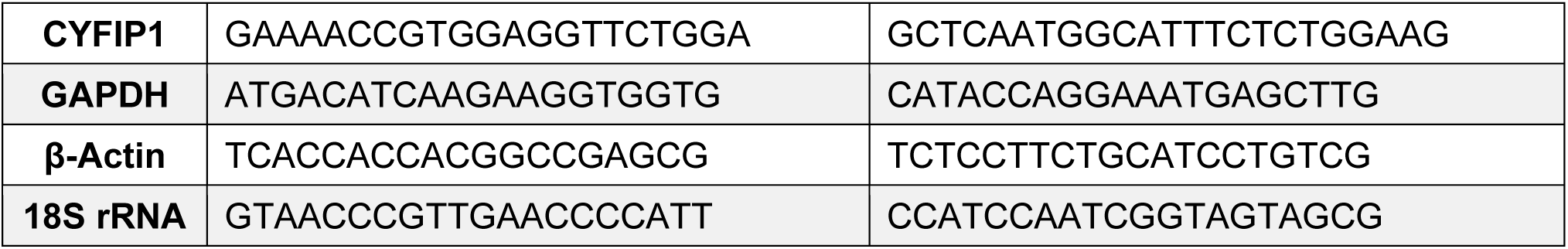

### RNA extraction and sequencing

Total RNA was extracted from TRIzol lysates using the PureLink RNA mini kit (Ambion, ThermoFisher). Three biological replicates were produced for each cell line and time-points of differentiation. RNA integrity was analysed using the RNA 6000 nano chip for the Agilent Bioanalyzer 2100 and only samples with a RIN > 9 were used for sequencing. The mRNA Hyperprep KAPA kit (Roche) was used for isolating mRNA and constructing the mRNA libraries. Indexed samples were pooled together and sequenced on an Illumina HiSeq 4000 (Illumina, San Diego, USA). Paired-end sequencing and a depth of approximately 33 million reads per sample were used for all sequencing experiments.

### Sterol extraction, derivatization and LC-MS

Isotope-labelled internal standards were purchased from Avanti Polar Lipids Inc (AL, USA), including 24R/S-hydroxycholesterol-D6, 7α-hydroxycholesterol-D7, 7-oxocholesterol-D7, 22S-3O-hydroxycholesterol-D7, 7α, 25-dihydroxycholsterol-D6, 3β-hydroxycholest-5-en-26-oic acid-D5, 7αH,3O-hydroxycholestenoic acid-D3, desmosterol-D6 and cholesterol-D7. Sterols were extracted from cell pellet through the dropwise addition of 1ml ethanol containing the internal standards under sonication in an ultrasonic bath. Samples were then centrifuged at 16,000g (13,000rpm) @ 4°C for 30mins. Supernatant was then collected and diluted with water to give a 70% ethanol solution. Oxysterols were separated from cholesterol by C18 solid-phase extraction and derivatized with Girard P reagent (US Biological) with or without prior oxidation of 3β-hydroxy to 3-oxo groups. The method for derivatization and LC-MS has been previously described in detail [3,4]. The identification of 24S,24-EC is based on the accurate mass of parent ions (≤5 ppm) and MS3 fragmentation spectrum which are matched to the authentic standard.

**Supplementary table 9.** Antibodies used for immunocytochemistry.

| Target | Species | Dilution | Cat. Number | Supplier |
| --- | --- | --- | --- | --- |
| **CTIP2** | Rat | 1:500 | AB18465 | Abcam |
| **FOXG1** | Rabbit | 1:1000 | AB18259 | Abcam (ChIP grade) |
| **GABA** | Rabbit | 1:1000 | A2052 | Sigma |
| **Ki67** | Mouse | 1:1000 | 550609 | BD Pharmingen |
| **NESTIN** | Rabbit | 1:500 | ABD69 | Millipore |
| **NeuN** | Mouse | 1:500 | MAB377 | Millipore |
| **NeuN** | Rabbit | 1:500 | ABN78 | Millipore |
| **OCT4** | Goat | 1:500 | SC-8629 | Santa Cruz |
| **P27<sup>KIP1</sup>** | Rabbit | 1:500 | 06-445 | Upstate (Millipore) |
| **PAX6** | Mouse | 1:1000 | PAX6 | DSHB |
| **PH3** | Rat | 1:1000 | 641002 | Biologend |
| **SATB2** | Mouse | 1:100 | AB51502 | Abcam |
| **SSEA3** | Rat | 1:500 | MAB4303 | Millipore |
| **TBR1** | Rabbit | 1:500 | AB31940 | Abcam |

**Supplementary table 10.** Antibodies used for Western Blot.

| Target | Species | Dilution | Cat. Number | Supplier |
| --- | --- | --- | --- | --- |
| **βCAT** | mouse | 1:1000 | Sc-7963 | Santa Cruz |
| **CYFIP1** | rabbit | 1:3000 | AB6046 | Millipore |
| **GAPDH** | mouse | 1:10000 | AB8245 | Abcam |
| **GAPDH** | rabbit | 1:10000 | AB9485 | Abcam |
| **NIPA1** | rabbit | 1:500 | AB128640 | Abcam |
| **NIPA2** | rabbit | 1:500 | AB84343 | Abcam |
| **TUBGCP5** | mouse | 1:1000 | AB168325 | Abcam |
| **LXRβ** | rabbit | 1:1000 | NB100-74457 | Novus Biologicals |

## References

1 Gejman, P. V., Sanders, A. R. & Kendler, K. S. Genetics of schizophrenia: new findings and challenges. Annu Rev Genomics Hum Genet 12, 121–144, doi:10.1146/annurev-genom-082410-101459 (2011).

2 International Schizophrenia, C. Rare chromosomal deletions and duplications increase risk of schizophrenia. Nature 455, 237–241, doi:10.1038/nature07239 (2008).

3 Kirov, G. et al. Support for the involvement of large copy number variants in the pathogenesis of schizophrenia. Hum Mol Genet 18, 1497–1503, doi:10.1093/hmg/ddp043 (2009).

4 Kirov, G. et al. The penetrance of copy number variations for schizophrenia and developmental delay. Biol Psychiatry 75, 378–385, doi:10.1016/j.biopsych.2013.07.022 (2014).

5 Stefansson, H. et al. Large recurrent microdeletions associated with schizophrenia. Nature 455, 232–236, doi:10.1038/nature07229 (2008).

6 Rees, E. et al. Analysis of copy number variations at 15 schizophrenia associated loci. Br J Psychiatry 204, 108–114, doi:10.1192/bjp.bp.113.131052 (2014).

7 Picinelli, C. et al. Recurrent 15q11.2 BP1-BP2 microdeletions and microduplications in the etiology of neurodevelopmental disorders. Am J Med Genet B Neuropsychiatr Genet 171, 1088–1098, doi:10.1002/ajmg.b.32480 (2016).

8 Chai, J. H. et al. Identification of four highly conserved genes between breakpoint hotspots BP1 and BP2 of the Prader-Willi/Angelman syndromes deletion region that have undergone evolutionary transposition mediated by flanking duplicons. Am J Hum Genet 73, 898–925, doi:S0002-9297(07)63637-4 [pii] 10.1086/378816 (2003).

9 Schenck, A., Bardoni, B., Moro, A., Bagni, C. & Mandel, J. L. A highly conserved protein family interacting with the fragile X mental retardation protein (FMRP) and displaying selective interactions with FMRP-related proteins FXR1P and FXR2P. Proc Natl Acad Sci U S A 98, 8844–8849, doi:10.1073/pnas.151231598151231598 [pii] (2001).

10 Napoli, I. et al. The fragile X syndrome protein represses activity-dependent translation through CYFIP1, a new 4E-BP. Cell 134, 1042–1054, doi:10.1016/j.cell.2008.07.031 S0092-8674(08)00951-3 [pii] (2008).

11 Chen, Z. et al. Structure and control of the actin regulatory WAVE complex. Nature 468, 533–538, doi:10.1038/nature09623 (2010).

12 Oguro-Ando, A. et al. Increased CYFIP1 dosage alters cellular and dendritic morphology and dysregulates mTOR. Mol Psychiatry 20, 1069–1078, doi:10.1038/mp.2014.124 mp2014124 [pii] (2015).

13 Pathania, M. et al. The autism and schizophrenia associated gene CYFIP1 is critical for the maintenance of dendritic complexity and the stabilization of mature spines. Transl Psychiatry 4, e374, doi:10.1038/tp.2014.16 tp201416 [pii] (2014).

14 Davenport, E. C. et al. Autism and Schizophrenia-Associated CYFIP1 Regulates the Balance of Synaptic Excitation and Inhibition. Cell Rep 26, 2037–2051 e2036, doi:10.1016/j.celrep.2019.01.092 (2019).

15 Sahasrabudhe, A. et al. Cyfip1 Regulates SynGAP1 at Hippocampal Synapses. Front Synaptic Neurosci 12, 581714, doi:10.3389/fnsyn.2020.581714 (2020).

16 Habela, C. W. et al. Persistent Cyfip1 Expression Is Required to Maintain the Adult Subventricular Zone Neurogenic Niche. J Neurosci 40, 2015–2024, doi:10.1523/JNEUROSCI.2249-19.2020 JNEUROSCI.2249-19.2020 [pii] (2020).

17 Haan, N. et al. Haploinsufficiency of the schizophrenia and autism risk gene Cyfip1 causes abnormal postnatal hippocampal neurogenesis through microglial and Arp2/3 mediated actin dependent mechanisms. Transl Psychiatry 11, 313, doi:10.1038/s41398-021-01415-6 (2021).

18 Dominguez-Iturza, N. et al. The autism- and schizophrenia-associated protein CYFIP1 regulates bilateral brain connectivity and behaviour. Nat Commun 10, 3454, doi:10.1038/s41467-019-11203-y (2019).

19 Silva, A. I. et al. Cyfip1 haploinsufficient rats show white matter changes, myelin thinning, abnormal oligodendrocytes and behavioural inflexibility. Nat Commun 10, 3455, doi:10.1038/s41467-019-11119-7 (2019).

20 Yoon, K. J. et al. Modeling a genetic risk for schizophrenia in iPSCs and mice reveals neural stem cell deficits associated with adherens junctions and polarity. Cell Stem Cell 15, 79–91, doi:10.1016/j.stem.2014.05.003S1934-5909(14)00190-8 [pii] (2014).

21 Stefansson, H. et al. CNVs conferring risk of autism or schizophrenia affect cognition in controls. Nature 505, 361–366, doi:10.1038/nature12818 (2014).

22 Gonzalez, F. et al. An iCRISPR platform for rapid, multiplexable, and inducible genome editing in human pluripotent stem cells. Cell Stem Cell 15, 215–226, doi:10.1016/j.stem.2014.05.018 (2014).

23 Nowakowski, T. J. et al. Spatiotemporal gene expression trajectories reveal developmental hierarchies of the human cortex. Science 358, 1318–1323, doi:10.1126/science.aap8809 (2017).

24 Nguyen, L. et al. p27kip1 independently promotes neuronal differentiation and migration in the cerebral cortex. Genes Dev 20, 1511–1524, doi:10.1101/gad.377106 (2006).

25 Jiao, Z. et al. Dopachrome tautomerase (Dct) regulates neural progenitor cell proliferation. Dev Biol 296, 396–408, doi:10.1016/j.ydbio.2006.06.006 (2006).

26 Greif, K. F., Asabere, N., Lutz, G. J. & Gallo, G. Synaptotagmin-1 promotes the formation of axonal filopodia and branches along the developing axons of forebrain neurons. Dev Neurobiol 73, 27–44, doi:10.1002/dneu.22033 (2013).

27 Sacchetti, P. et al. Liver X receptors and oxysterols promote ventral midbrain neurogenesis in vivo and in human embryonic stem cells. Cell Stem Cell 5, 409–419, doi:10.1016/j.stem.2009.08.019 S1934-5909(09)00403-2 [pii] (2009).

28 Theofilopoulos, S. et al. 24(S),25-Epoxycholesterol and cholesterol 24S hydroxylase (CYP46A1) overexpression promote midbrain dopaminergic neurogenesis in vivo. J Biol Chem 294, 4169–4176, doi:10.1074/jbc.RA118.005639 S0021-9258(20)41824-1 [pii] (2019).

29 Theofilopoulos, S. et al. Brain endogenous liver X receptor ligands selectively promote midbrain neurogenesis. Nat Chem Biol 9, 126–133, doi:10.1038/nchembio.1156 nchembio.1156 [pii] (2013).

30 Nowakowski, R. S., Lewin, S. B. & Miller, M. W. Bromodeoxyuridine immunohistochemical determination of the lengths of the cell cycle and the DNA-synthetic phase for an anatomically defined population. J Neurocytol 18, 311–318, doi:10.1007/BF01190834 (1989).

31 Lehmann, J. M. et al. Activation of the nuclear receptor LXR by oxysterols defines a new hormone response pathway. J Biol Chem 272, 3137–3140, doi:10.1074/jbc.272.6.3137 S0021-9258(19)78355-0 [pii] (1997).

32 Janowski, B. A. et al. Structural requirements of ligands for the oxysterol liver X receptors LXRalpha and LXRbeta. Proc Natl Acad Sci U S A 96, 266–271, doi:10.1073/pnas.96.1.266 (1999).

33 Willy, P. J. et al. LXR, a nuclear receptor that defines a distinct retinoid response pathway. Genes Dev 9, 1033–1045, doi:10.1101/gad.9.9.1033 (1995).

34 Pehkonen, P. et al. Genome-wide landscape of liver X receptor chromatin binding and gene regulation in human macrophages. BMC Genomics 13, 50, doi:10.1186/1471-2164-13-50 1471-2164-13-50 [pii] (2012).

35 Savory, J. G., Edey, C., Hess, B., Mears, A. J. & Lohnes, D. Identification of novel retinoic acid target genes. Dev Biol 395, 199–208, doi:10.1016/j.ydbio.2014.09.013 (2014).

36 Durak, O. et al. Chd8 mediates cortical neurogenesis via transcriptional regulation of cell cycle and Wnt signaling. Nat Neurosci 19, 1477–1488, doi:10.1038/nn.4400 (2016).

37 Mariani, J. et al. FOXG1-Dependent Dysregulation of GABA/Glutamate Neuron Differentiation in Autism Spectrum Disorders. Cell 162, 375–390, doi:10.1016/j.cell.2015.06.034 (2015).

38 Marchetto, M. C. et al. Altered proliferation and networks in neural cells derived from idiopathic autistic individuals. Mol Psychiatry 22, 820–835, doi:10.1038/mp.2016.95 (2017).

39 Paulsen, B. et al. Autism genes converge on asynchronous development of shared neuron classes. Nature 602, 268–273, doi:10.1038/s41586-021-04358-6 (2022).

40 Kim, N. S. et al. CYFIP1 Dosages Exhibit Divergent Behavioral Impact via Diametric Regulation of NMDA Receptor Complex Translation in Mouse Models of Psychiatric Disorders. Biol Psychiatry 92, 815–826, doi:10.1016/j.biopsych.2021.04.023 (2022).

41 Silva, A. I. et al. Analysis of Diffusion Tensor Imaging Data From the UK Biobank Confirms Dosage Effect of 15q11.2 Copy Number Variation on White Matter and Shows Association With Cognition. Biol Psychiatry 90, 307–316, doi:10.1016/j.biopsych.2021.02.969 (2021).

42 Kirkeby, A. et al. Generation of regionally specified neural progenitors and functional neurons from human embryonic stem cells under defined conditions. Cell Rep 1, 703–714, doi:10.1016/j.celrep.2012.04.009 (2012).

43 Wang, Y. et al. Targeted lipidomic analysis of oxysterols in the embryonic central nervous system. Mol Biosyst 5, 529–541, doi:10.1039/b819502a (2009).

44 Wong, J., Quinn, C. M., Guillemin, G. & Brown, A. J. Primary human astrocytes produce 24(S),25-epoxycholesterol with implications for brain cholesterol homeostasis. J Neurochem 103, 1764–1773, doi:10.1111/j.1471-4159.2007.04913.x (2007).

45 Arai, Y. et al. Neural stem and progenitor cells shorten S-phase on commitment to neuron production. Nat Commun 2, 154, doi:10.1038/ncomms1155 (2011).

46 Parente, M. et al. Brain Cholesterol Biosynthetic Pathway Is Altered in a Preclinical Model of Fragile X Syndrome. Int J Mol Sci 23, doi:10.3390/ijms23063408 (2022).

47 Tomita, H. et al. 7-Dehydrocholesterol-derived oxysterols cause neurogenic defects in Smith-Lemli-Opitz syndrome. Elife 11, doi:10.7554/eLife.67141 (2022).

48 Thurm, A. et al. Development, behavior, and biomarker characterization of Smith-Lemli-Opitz syndrome: an update. J Neurodev Disord 8, 12, doi:10.1186/s11689-016-9145-x (2016).

49 Leoni, V. & Caccia, C. Oxysterols as biomarkers in neurodegenerative diseases. Chem Phys Lipids 164, 515–524, doi:10.1016/j.chemphyslip.2011.04.002 (2011).

50 Bjorkhem, I. Crossing the barrier: oxysterols as cholesterol transporters and metabolic modulators in the brain. J Intern Med 260, 493–508, doi:JIM1725 [pii] 10.1111/j.1365-2796.2006.01725.x (2006).

51 Gamba, P. et al. The link between altered cholesterol metabolism and Alzheimer’s disease. Ann N Y Acad Sci 1259, 54–64, doi:10.1111/j.1749-6632.2012.06513.x (2012).

52 Karasinska, J. M. & Hayden, M. R. Cholesterol metabolism in Huntington disease. Nat Rev Neurol 7, 561–572, doi:10.1038/nrneurol.2011.132 nrneurol.2011.132 [pii] (2011).

53 Jin, U., Park, S. J. & Park, S. M. Cholesterol Metabolism in the Brain and Its Association with Parkinson’s Disease. Exp Neurobiol 28, 554–567, doi:10.5607/en.2019.28.5.554 en.2019.28.5.554 [pii] (2019).

54 Sun, Z. et al. Brain-Specific Oxysterols and Risk of Schizophrenia in Clinical High-Risk Subjects and Patients With Schizophrenia. Front Psychiatry 12, 711734, doi:10.3389/fpsyt.2021.711734 (2021).

55 Grayaa, S. et al. Plasma oxysterol profiling in children reveals 24 hydroxycholesterol as a potential marker for Autism Spectrum Disorders. Biochimie 153, 80–85, doi:10.1016/j.biochi.2018.04.026 (2018).

56 Guidara, W. et al. Plasma oxysterols: Altered level of plasma 24 hydroxycholesterol in patients with bipolar disorder. J Steroid Biochem Mol Biol 211, 105902, doi:10.1016/j.jsbmb.2021.105902 (2021).

57 Fernandez Cardo, L., de la Fuente, D. C. and Li, M. Impaired neurogenesis and neural progenitor fate choice in a human stem cell model of SETBP1 disorder. Molecular Autism 14, doi:https://doi.org/10.1186/s13229-023-00540-x (2023).

58 Writing Committee for the, E.-C. N. V. W. G., et al. Association of Copy Number Variation of the 15q11.2 BP1-BP2 Region With Cortical and Subcortical Morphology and Cognition. JAMA Psychiatry 77, 420–430, doi:10.1001/jamapsychiatry.2019.3779 (2020).

59 Brown, M. S. & Goldstein, J. L. The SREBP pathway: regulation of cholesterol metabolism by proteolysis of a membrane-bound transcription factor. Cell 89, 331–340, doi:10.1016/s0092-8674(00)80213-5 (1997).

60 Sakakura, Y. et al. Sterol regulatory element-binding proteins induce an entire pathway of cholesterol synthesis. Biochem Biophys Res Commun 286, 176–183, doi:10.1006/bbrc.2001.5375 (2001).

61 Darnell, J. C. et al. FMRP stalls ribosomal translocation on mRNAs linked to synaptic function and autism. Cell 146, 247–261, doi:10.1016/j.cell.2011.06.013 (2011).

62 Sawicka, K. et al. FMRP has a cell-type-specific role in CA1 pyramidal neurons to regulate autism-related transcripts and circadian memory. Elife 8, doi:10.7554/eLife.46919 (2019).

63 Ascano, M., Jr., et al. FMRP targets distinct mRNA sequence elements to regulate protein expression. Nature 492, 382–386, doi:10.1038/nature11737 (2012).

64 Sperandeo, A. et al. Cortical neuronal hyperexcitability and synaptic changes in SGCE mutation-positive myoclonus dystonia. Brain, doi:10.1093/brain/awac365 (2022).

65 Dobin, A. et al. STAR: ultrafast universal RNA-seq aligner. Bioinformatics 29, 15–21, doi:10.1093/bioinformatics/bts635 (2013).

66 Yu, G., Wang, L. G., Han, Y. & He, Q. Y. clusterProfiler: an R package for comparing biological themes among gene clusters. OMICS 16, 284–287, doi:10.1089/omi.2011.0118 (2012).

## References

1. Zhao, S.; Maxwell, S.; Jimenez-Beristain, A.; Vives, J.; Kuehner, E.; Zhao, J.; O’Brien, C.; de Felipe, C.; Semina, E.; Li, M. Generation of embryonic stem cells and transgenic mice expressing green fluorescence protein in midbrain dopaminergic neurons. Eur J Neurosci 2004, 19, 1133–1140, doi:10.1111/j.1460-9568.2004.03206.x.

2. Gonzalez, F.; Zhu, Z.; Shi, Z.D.; Lelli, K.; Verma, N.; Li, Q.V.; Huangfu, D. An iCRISPR platform for rapid, multiplexable, and inducible genome editing in human pluripotent stem cells. Cell Stem Cell 2014, 15, 215–226, doi:10.1016/j.stem.2014.05.018.

3. Crick, P.J.; William Bentley, T.; Abdel-Khalik, J.; Matthews, I.; Clayton, P.T.; Morris, A.A.; Bigger, B.W.; Zerbinati, C.; Tritapepe, L.; Iuliano, L.;, et al. Quantitative charge-tags for sterol and oxysterol analysis. Clin Chem 2015, 61, 400–411, doi:10.1373/clinchem.2014.231332.

4. Yutuc, E.; Dickson, A.L.; Pacciarini, M.; Griffiths, L.; Baker, P.R.S.; Connell, L.; Ohman, A.; Forsgren, L.; Trupp, M.; Vilarinho, S.;, et al. Deep mining of oxysterols and cholestenoic acids in human plasma and cerebrospinal fluid: Quantification using isotope dilution mass spectrometry. Anal Chim Acta 2021, 1154, 338259, doi:10.1016/j.aca.2021.338259.

